# Inhibition of miR-1307 Reverses Resistance to Cisplatin in Drug-Resistant Oral Squamous Cell Carcinoma

**DOI:** 10.64898/2026.04.06.709730

**Authors:** Aditi Patel, Vaishnavi Patel, Shreya Lotia, Kaustubh Patel, Dushyant Mandlik, Jonathan Tan, Prabha Sampath, Binita Patel, Kaid Johar, Dhiraj Bhatia, Vivek Tanavde, Shanaya Patel

**Author notes:** **Corresponding author details:** Shanaya Patel, PhD, DBT/Wellcome Trust Early Career Fellow, Division of Biological & Life Sciences, School of Arts & Sciences, Ahmedabad University, Central Campus, Navrangpura, Ahmedabad, Gujarat-380009, India., Vivek Tanavde, PhD, Associate Dean Sciences, Associate Professor, Division of Biological & Life Sciences, School of Arts & Sciences, Ahmedabad University, Central Campus, Navrangpura, Ahmedabad, Gujarat-380009, India.

## Abstract

**Background:** Chemo-resistance remains a major clinical challenge in Oral Squamous Cell Carcinoma (OSCC), attributed to the intrinsically resistant cells. Although tumour-derived extracellular vesicles (EVs) have been implicated in cell–cell communication, their role in propagating chemo-resistance remains poorly defined. This study aims to identify salivary EV-associated miRNAs capable of predicting chemoresistance and to delineate the role of miR-1307-5p in modulating CSC-driven therapeutic refractoriness.

**Methods:** Salivary EV-derived expression profile of miR-1307-5p was assessed by qPCR in chemo resistant OSCC patients and further validated in TCGA small RNA sequencing datasets. Expression was validated by qPCR and correlated with clinicopathological outcomes. Functional assays including cell-cycle analysis, apoptosis, migration/invasion, 3D spheroids, angiogenesis, and CAM assays were performed in miR-1307-5p inhibited CD44⁺ CSC subpopulation compared to its vehicular control. Transcriptomic profiling cross-referencing with TCGA was conducted to identify potential novel targets of miR-1307-5p. Chemo-sensitisation was assessed by treating the knockdown chemo resistant cells with low dose cisplatin and validating it using *in-vitro* functional assays and orthotopic xenograft model.

**Results:** miR-1307-5p was significantly elevated in salivary EVs of chemo resistant OSCC patients and correlated with poor overall survival (p = 0.03). The miRNA was markedly enriched in endogenously resistant CD44⁺ CSCs. Silencing of miR-1307-5p induced G2/M arrest, triggered apoptosis, impaired invasion, and reduced angiogenesis both *in-vitro* and in *ex-vivo* assays. Transcriptomic profiling, TCGA validation, and integrative pathway analysis identified key oncogenic hubs which converge on PI3K–AKT, MAPK/ERK, and YAP signalling pathways governing EMT. Inhibition of miR-1307-5p restored cisplatin sensitivity in resistant CSCs, with low-dose cisplatin producing substantial tumour suppression *in-vitro* and *in-vivo*. Reduced CD44 expression in xenograft models confirmed CSC reprogramming. EVs from anti-miR-treated cells confer chemo sensitisation upon uptake by resistant CSCs. Xenograft models substantiated that EVs can initiate tumour formation and that EV-mediated delivery of anti-miR-1307-5p drives significant tumour regression.

**Conclusion:** This study identifies salivary EV-derived miR-1307-5p as a clinically relevant biomarker of chemoresistance in OSCC and reveals its mechanistic role in sustaining CSC-driven therapeutic failure. Targeting miR-1307-5p offers a promising avenue for restoring cisplatin sensitivity and developing exosome-based therapeutic strategies.

**Graphical Abstract:** 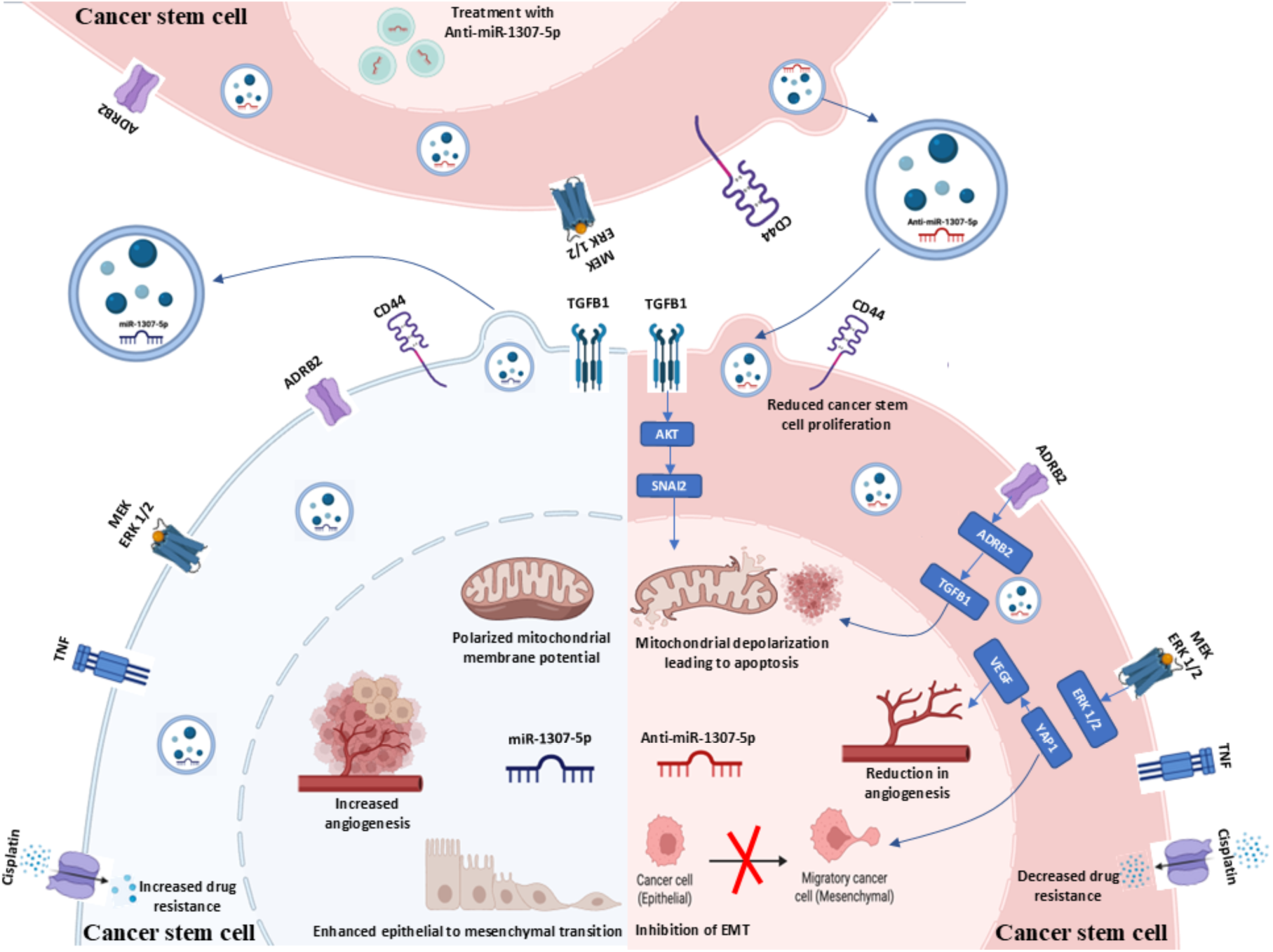

## Background

Oral squamous cell carcinoma (OSCC) remains a major global health burden, with GLOBOCAN 2022 reporting over 3,89,846 new cases worldwide (1). Despite multimodal therapy, a substantial proportion of patients require neo-adjuvant or adjuvant chemotherapy, with 30–40% exhibiting resistance to standard chemotherapeutic regimens (2). Platinum-based drugs remain the gold standard of OSCC therapy; however, patients with platinum-refractory disease experience a median overall survival of <6 months, indicating the urgent need for improved therapeutic strategies and a deeper understanding of resistance mechanisms (3). Hence, there is a clinical need for minimally invasive biomarkers capable of predicting chemoresistance preceding clinical progression, that can enable timely therapeutic intervention.

Among liquid biopsy platforms, saliva has gained significant attention in OSCC due to its accessibility, non-invasiveness, and tumour proximity (4). Salivary microRNAs (miRNAs), known regulators of nearly 60% of protein-coding genes and many CSC-associated pathways, offer strong prognostic potential (5). However, free miRNAs in saliva are susceptible to degradation by nucleases, limiting their clinical utility. Extracellular vesicles (EVs) overcome this limitation by protecting miRNAs within a lipid bilayer and facilitating their stable detection (6). EV-associated miRNAs actively mediate intercellular communication and can transfer oncogenic traits, contributing to tumour progression, pre-metastatic niche formation, and drug resistance (6). Although EV-miRNAs have been linked to therapeutic response in several cancers, their role in chemoresistance of OSCC is minimally explored. Existing OSCC-related miRNA biomarkers (e.g., miR-21, miR-491-5p, miR-196a (7-9)) are limited in their diagnostic or prognostic utility. However, no salivary EV-miRNA biomarkers predicting chemoresistance have been reported in OSCC.

This study aims to evaluate the clinical utility of miR-1307-5p in predicting chemotherapeutic resistance and to elucidate the role of miR-1307-5p in driving sensitive cells to a resistant phenotype. The insights from this study will highlight the utility of EV-derived miR-1307-5p as a potential biomarker for patient stratification and treatment optimisation.

## Methods

### Sampling details

Samples of resected tumour tissue and unstimulated saliva were collected from patients diagnosed with OSCC (primary subsites: buccal mucosa, floor of mouth, gum, and palate) from HCG Cancer Centre, Ahmedabad. The study was approved by the Institutional Ethics Committee of HCG Cancer Centre (ECR/92lInst/GJ/2013/RR-19) and written informed consent was obtained from all patients. This study complied with the guidelines set forth by the Declaration of Helsinki (2008). All patients enrolled in this study had a history of smokeless tobacco consumption and had been administered cisplatin or a combination of cisplatin with 5-Fluorouracil and docetaxel as a neoadjuvant/ adjuvant chemotherapy. Stage I–IV OSCC patients with or without radiotherapy were also included. These patients were further categorised as chemo-responders (n=33) with complete pathological responses for ≥3 years or chemo-non-responders (n=52) with stable/progressive disease post-treatment, per Response Evaluation Criteria in Solid Tumours (RECIST 1.1) criteria (Table 3.1). Power analysis was performed to estimate the minimum sample size required to detect differences in miR-1307-5p expression between chemo-responders and non-responders. Patients with HIV/HbsAg/HPV infections, leukoplakia lesions, paediatric cases, or OSCC-related complications were excluded. All sample processing, clinical evaluation, and statistical analysis were performed in a fully blinded manner to minimise bias. Unstimulated whole saliva of healthy individuals (matched for age and gender with the patients, n=33) with no etiological history of tobacco chewing and no clinically detectable oral lesions were used as normal controls.

**Table 1-.**
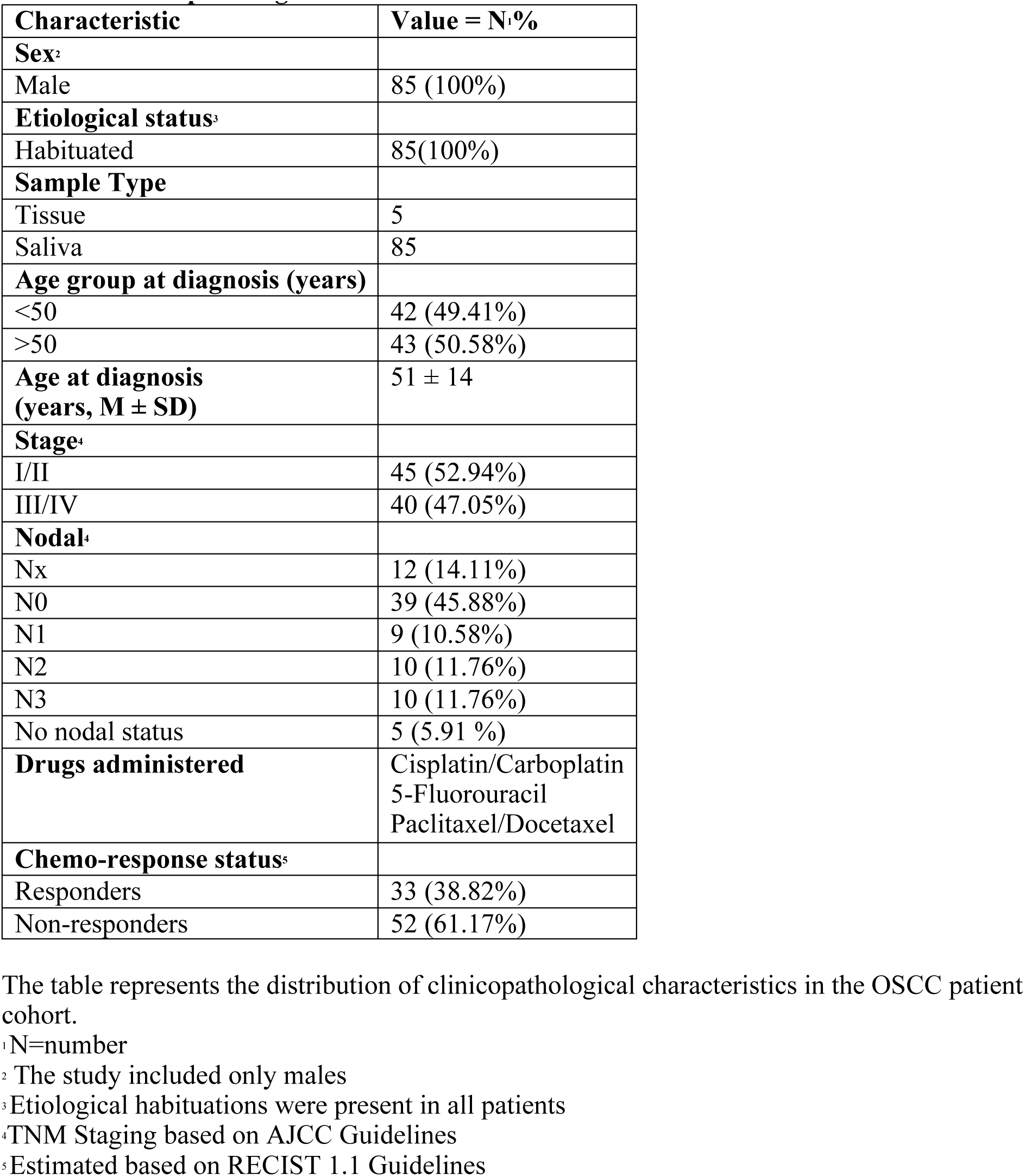
Clinicopathological characteristics of the OSCC patient cohort.

| **Characteristic** | **Value = N: %** |
| --- | --- |
| **Sex<sup>2</sup>** |  |
| Male | 85 (100%) |
| **Etiological status<sup>3</sup>** |  |
| Habituated | 85(100%) |
| **Sample Type** |  |
| Tissue | 5 |
| Saliva | 85 |
| **Age group at diagnosis (years)** |  |
| <50 | 42 (49.41%) |
| >50 | 43 (50.58%) |
| **Age at diagnosis (years, M ± SD)** | 51 ± 14 |
| **Stage<sup>4</sup>** |  |
| I/II | 45 (52.94%) |
| III/IV | 40 (47.05%) |
| **Nodal<sup>4</sup>** |  |
| Nx | 12 (14.11%) |
| N0 | 39 (45.88%) |
| N1 | 9 (10.58%) |
| N2 | 10 (11.76%) |
| N3 | 10 (11.76%) |
| No nodal status | 5 (5.91 %) |
| **Drugs administered** | Cisplatin/Carboplatin<br>5-Fluorouracil<br>Paclitaxel/Docetaxel |
| **Chemo-response status<sup>5</sup>** |  |
| Responders | 33 (38.82%) |
| Non-responders | 52 (61.17%) |

The table represents the distribution of clinicopathological characteristics in the OSCC patient cohort.
<sup>1</sup> N=number
<sup>2</sup> The study included only males
<sup>3</sup> Etiological habituations were present in all patients
<sup>4</sup> TNM Staging based on AJCC Guidelines
<sup>5</sup> Estimated based on RECIST 1.1 Guidelines

### EV isolation and characterisation

1-2 ml of whole saliva was collected from patient and control cohorts in a sterile tube and centrifuged at 2000 × g for 10 minutes at room temperature. EVs were isolated using the Total EVs Isolation Reagent from Other Body Fluids kit (Thermo Fisher Scientific, USA) according to the manufacturer’s protocol. For isolation of EVs from cell culture supernatant was centrifuged at 2000 × g for 10 minutes at 37°C and then processed using Total Exosome Isolation Reagent (Thermo Fisher Scientific, USA) followed by overnight incubation at 4°C according to manufacturer’s protocol. Post centrifugation EVs were eluted in 50 µL PBS. EV characterisation was carried out according to the previously described protocol (10). The size and concentration of the EVs were determined using Nanoparticle Tracking Analysis (NTA) on NanoSight LM10 (Malvern Panalytical Ltd., UK), and the morphology was assessed using Jeol JEM 1010 transmission electron microscope (Jeol, Japan). Expression of EV-tetraspanins was determined using PE-tagged antibodies-CD9, CD63, and CD81 (Cellarcus Biosciences, USA) and FITC-tagged CD47 (Becton Dickinson Biosciences, USA) on the CytoFLEX-LX (Beckman Coulter, USA) flow cytometer as mentioned previously (10).

### miRNA *in-situ* hybridisation

Formalin-fixed, paraffin-embedded (FFPE) tissue sections (5 µm) from 10 OSCC patients were subjected to heat-induced epitope retrieval using a pretreatment solution (Panomics, USA), followed by washing with PBS and protease digestion at 37 °C (Panomics, USA). The sections were then incubated with 5′ DIG-labelled LNA probes specific for miR-1307-5p (Exiqon, Denmark) in a hybridisation buffer (Roche, Switzerland) at 51 °C for 4 hours. After blocking with 10% goat serum, sections were incubated with anti-DIG alkaline phosphatase antibody (RRID: AB_514497) (Roche, Switzerland). Signal detection was performed using Fast Red substrate (Panomics, USA), followed by DAPI counterstaining. Finally, sections were mounted with FluorSave™ (Sigma-Aldrich, Merck, USA), and images were acquired using an Olympus FluoView FV1000 microscope equipped with a TRITC filter (Olympus, Japan).

### Cell culture and CD44+ cells isolation

Human Oral Squamous Cell Carcinoma cell line (OECM-1) was obtained from Sigma Aldrich (Merck, USA). RPMI1640 medium (Thermo Fisher Scientific, USA) supplemented with 10% Fetal bovine serum (FBS) (Himedia, India) and 1% Antibiotics and Antimycotics was used to maintain the cell line (Thermo Fisher Scientific, USA) at 37°C and 5% CO2. CD44+ CSCs were isolated using CD44 microbead-labelled cells on MACS columns (Miltenyi Biotec, Germany). The characterisation of the CD44+ population was characterised using flow cytometric analysis. The cells were stained, were suspended in 1 ml 1X PBS and stained with 5 µl of PE anti-human CD44 antibody (Miltenyi Biotec, USA) for 20 minutes in dark, washed with 1X PBS and resuspended in 100 µl 1X PBS. The acquisition of cells was performed on CytoFLEX Flow Cytometer (Beckman Coulter, USA), and the CD44+ population was determined by plotting a dot plot of B610-A (Phycoerythrin Area) vs SSC (Side scatter).

### Cell Treatments

CD44+ CSCs were transfected with miR-1307-5p inhibitor (anti-miR-1307-5p) or scrambled control (SC) (Qiagen, Germany) at initial concentrations ranging from 2.5 nM to 75 nM using Lipofectamine^TM^ 2000 reagent (Invitrogen, USA) for 48 hours.

To evaluate the chemo-reversal effect in CD44+ CSCs transfected with anti-miR-1307-5p or SC at their respective IC50 values, they were subsequently exposed to cisplatin (Sigma, USA) at varying concentrations (0.01–10 µM) for 24 hours.

To determine whether EVs mediate the transfer of chemo-resistance–modulating signals, EVs were isolated from anti-miR-1307-5p and cisplatin-treated CD44⁺ CSCs following 12 hours of starvation in serum-free media. Functional analyses were performed by co-culturing CD44+ CSCs with isolated EVs at an initial concentration ranging from 3 × 10^6^ particles/mL to 15.4 × 10^6^ particles/mL.

### Evaluation of EV Internalisation

CD44+ CSCs were incubated with 10 µg/mL of DiL-labeled EVs (1,1’-Dioctadecyl-3,3,3’,3’-Tetramethylindocarbocyanine Perchlorate) in culture medium for 3 hours at 37°C. Post PBS washes cells were fixed with 4% paraformaldehyde at room temperature for 15 minutes. Nuclei were counterstained with DAPI, and samples were mounted for imaging. EV internalisation was analysed using Confocal microscopy, (Leica DMI8, Germany).

### Quantitative Real-time PCR (qPCR) analysis

RNA isolation from salivary EVs, and supernatant-derived EVs and CD44+ CSCs was carried out using TRIzol^TM^ reagent (Thermo Fisher Scientific, USA) as per the previously mentioned protocol [10]. Post ethanol washes and air-drying the RNA pellet was resuspended in 25 µl nuclease-free water and quantified using Qubit™ Fluorometer 4.0 (Thermo Fisher Scientific, USA). Complementary DNA (cDNA) conversion was carried out using miRNA-specific miRCury LNA RT kit (QIAGEN, Germany) (cycling conditions: 42°C for 60 minutes, followed by 95°C for 5 minutes) and gene-specific QuantiTect® Reverse Transcription Kit (QIAGEN, Germany). Further, qPCR was performed using SYBR Fast Universal Mix (KAPA, Sigma-Aldrich, USA) on QuantStudio^TM^ 5 Real-Time PCR (Thermo Fischer Scientific, USA) (cycling conditions: 95°C for 3 minutes, 95°C for 3 seconds followed by 60°C for 20 seconds). U6 and β-actin were used as internal controls, and fold change was calculated using the 2−ΔΔCt method. The primer sequences are listed in Supplementary Table 1.

### Cell Proliferation assay

The cytotoxicity induced by transfection, cisplatin and EV-based treatments on CD44+ cells was assessed using MTT (3-(4,5-dimethylthiazol-2-yl)-2,5-diphenyltetrazolium bromide) assay. 3000 cells/well were treated followed by addition of 10 µl (5mg/mL) of MTT. Dimethyl sulfoxide (DMSO) was used to solubilise the formazan crystals and proliferation was determined by measuring absorbance at 570 nm using a Synergy™ H1 microplate reader (Agilent BioTek Instruments, USA).

### Cell cycle assay

0.25 × 10^6^ cells/well post-treatment were harvested and fixed using 1 mL 70% chilled ethanol. Cells were incubated with 1 mL of 50 µg/mL PI (Propidium Iodide) solution (Becton Dickinson Biosciences, USA) for 30 minutes at 4°C in the dark and were acquired using the yellow-green laser (525 nm) on CytoFLEX-LX Flow Cytometer (Beckman Coulter Life Sciences, USA).

### Apoptosis assay

Annexin V-FITC Apoptosis Assay kit (Thermo Fisher Scientific, USA), was used to measure apoptosis. The harvested cells (1 × 10^6^ cells/mL) were resuspended in the 1X binding buffer, stained with Annexin V-FITC and PI for 10 minutes in dark and subjected to flow-cytometric analysis using CytoFLEX-LX Flow Cytometer (Beckman Coulter Life Sciences, USA).

### Nuclear Fragmentation assay

0.25 × 10^6^ cells were washed with PBS, fixed with 3:1 Methanol: Acetic acid fixative solution, followed by staining with PI dye (50 µg/mL) in the dark at 37°C. These cells were observed under the ZOE™ Fluorescent Cell Imager at 690 nm (BioRad laboratories, USA) and analysed by Fiji ImageJ (RRID:SCR_003070) software.

### Mitochondrial membrane potential estimation

The alteration in the mitochondrial membrane potential was determined using fluorescent dye 5,5,6,6’-tetrachloro-1,1’,3,3’ tetraethylbenzimi-dazoylcarbocyanine iodide (JC-1) (2mg/mL). 0.25 × 10^6^ cells/well were stained, incubated for 20 minutes in the dark and visualised under the ZOE™ Fluorescent Cell Imager (BioRad laboratories, USA). Changes in the mitochondrial membrane potential were estimated using the ratio of green to red fluorescence using Fiji ImageJ (RRID:SCR_003070) software.

### Analysis of Reactive Oxygen Species (ROS)

Intracellular ROS levels following treatment were quantified using 2′,7′-dichlorodihydrofluorescein diacetate (DCF-DA). 5 × 10³ cells per well were seeded, stained with DCF-DA (20 µM), and fluorescence intensity was measured at 488/525 nm using a Synergy™ H1 multimode plate reader (Agilent BioTek Instruments, USA).

### Wound Healing Assay

0.5 × 10^5^ cells/well were plated in a 24-well plate and a scratch was introduced using a 200 µL pipette tip. The wound closure was monitored for 48-hours using a 4X Bright-field microscope (Olympus microscope, Japan). The distance between the wounds was measured using the Fiji ImageJ (RRID:SCR_003070) software.

### Transwell Invasion Assay

The invasive ability of CD44+ CSCs post-treatment was determined using QCM^TM^ Collagen Cell Invasion Assay kit (Merck, Sigma-Aldrich, USA). Cells were serum starved for 8 hours and 0.5 × 10^6^ cells were harvested, placed in the upper chamber of the transwell and FBS-containing media was added to the lower chamber. Post incubation for 48 hours at 37°C, a detachment buffer was added, the inserts were removed, and CyQuant GR DyeR1 and 4X Cell Lysis Buffer were added followed by incubation in the dark. Fluorescence was measured at 480 nm/520 nm in a multimode plate reader (Synergy™ H1; Agilent BioTek Instruments, USA).

### 3D in-vitro hanging drop spheroid assay

Single-cell suspension of CD44+ cells was plated (5000 cells/mL) in a sterile petri-dish and incubated at 37 °C. Post 24 hours, spheroids were transferred to collagen-treated coverslips. The spheroids were fixed using 4% paraformaldehyde (PFA), stained with 0.1% phalloidin A488, and counterstained with DAPI. Spheroids were mounted using Mowiol and imaged using a confocal microscope (Leica DMI8, Germany) at 405 nm and 488 nm excitation/emission. The invasive ability of the spheroids was calculated with the Fiji ImageJ (RRID:SCR_003070) software using the formula: Cell invasion index = Migration distance of the cells from the surface of the spheroids/Diameter of the core of the spheroids

### Endothelial tube formation assay

The *in vitro* tube formation assay was conducted using Endothelial Cell Tube Formation Assay kit according to the manufacturer’s protocol (Thermo Fisher Scientific, USA). Human Large Vessel Endothelial Cells (HUVECs; 5 × 10^4^) were co-cultured with treated CD44+ CSCs and loaded onto the GeltrexTM-coated surface, supplemented with M200 medium. Capillary tube formation was monitored for 12 hours and then stained with Calcein (2 µg/mL) for 30 minutes at 37°C in the dark. Tube formation was visualised under 20X ZOE™ Fluorescent Cell Imager (BioRad Laboratories, USA) at 495-570 nm wavelengths and the branching points were quantified in three random fields using Fiji ImageJ (RRID:SCR_003070) software.

### Chick embryo chorioallantoic membrane (CAM) assay

Fertilised and pathogen-free eggs were sterilised using 70% isopropanol and incubated at 37°C with 60% humidity for eight days. Post-incubation, the eggs were dissected, injected with 5 × 10^4^ treated CD44+ cells, sealed and incubated for 48 hours at 37°C. The blood vessel development was monitored for 48 hours, and images were captured using a Nikon digital camera (Tokyo, Japan), The branching points of blood vessels were quantified using IKOSA software (KML Vision GmbH, Switzerland) for three random fields of each sample (11).

### Transcriptome sequencing and analysis

RNA was extracted from CD44+ control, SC, and anti-miR-1307-5p treated cells, and the library was prepared using NEBNext® Ultra^TM^ II RNA library prep kit Genotypic Technologies. Following quantification and RNA integrity assessment, Illumina HiSeq 2500 platform (150×2 chemistry) was used for sequencing (10, 12). Raw reads were processed with Trimmomatic (RRID:SCR_011848) (v0.32) [14], quality assessed with FastQC (RRID:SCR_014583) (v0.11.8) (13), and aligned to the human reference genome (GRCh38) using STAR (RRID:SCR_004463) (v2.11.8) (14). Transcript quantification and differential expression analysis were performed using StringTie (RRID SCR_016323) (v2.2.1) (15) and DESeq (RRID:SCR_000154) (v3.17) in R Bioconductor (RRID:SCR_006442) (16). Only transcripts with ≥40 read counts, log2FC≥2, and adjusted p-value<0.05 were considered significant.

### Analysis of The Cancer Genome Atlas (TCGA) datasets

miRNA and transcriptome sequencing data were retrieved from TCGA datasets (n=114) for HNSCC patients (subsites-Floor of mouth, Gum, Palate, and Other and unspecified parts of mouth). Further, 83 datasets were retrieved based on chemotherapy administration, and the patients were classified into two cohorts-chemo-responders and chemo-non-responders. The differential expression of the genes and miRNAs from these datasets was calculated using DESeq (RRID:SCR_000154) (v3.17). The miRNAs and gene expression were considered differentially expressed with a log2FC≥ +2 and ≤-2 and p-value< 0.05.

### Lentiviral vector construction

Lentiviral shRNA particles with the customised target sequence of sh-miR-1307-5p were purchased from Sigma Aldrich (Merck, USA) and the empty vectors were used as an internal control. 2 x 10^4^ CD44+ cells were seeded in a 96-well plate. Post 24 hours of incubation, the cells were treated with sh-miR-1307-5p viral particles with the Multiplicity of Infection (MOI)=2 and supplemented with media containing hexadimethrine bromide (8 µg/mL) for 24 hours. Stable shRNA vector constructs were chosen based on the puromycin kill curve (2500 ng/mL), and the selected cells were cultured for 2-3 weeks to establish stable cell lines.

### NOD-SCID mouse xenograft tumour model

5-9 weeks old, male NOD-SCID mice, weighing 20-25 grams (RRID:IMSR_JAX:001303) were obtained from the Indian Institute of Science Education and Research (IISER) Pune National Facility and were maintained at 23 ± 2°C humidity according to the protocol approved by Ethics Committee (IISER-PUNE/IAEC/2023_03/05). Mice were divided into four groups: negative control (PBS treated), positive control (OECM1 treated), Scrambled control (SC), and anti-miR-1307-5p transduced cells. The mice were anaesthetised with ketamine solution and injected with 1.5×10^6^ cells/ml into the oral cavity. For EV-based mice models, 3.5 x 10^5^ particles/mL were injected into the buccal cavity. Mice were monitored weekly for their body weight. Further, to understand the role of miR-1307-5p in chemo-reversal, mice with palpable tumours were administered 2 mg/kg cisplatin every 7th day for 2 weeks. 30 days post-injection, the mice were euthanised, and the tumour tissues and organs were resected and placed in 10% NBF (neutral buffered formalin) and 4% PFA for further analysis.

### Haemotoxylin and Eosin (H&E) Staining

The 4 µm tissue sections were dewaxed using xylene, rehydrated in absolute ethanol, 90% ethanol and 80% ethanol. Followed by haematoxylin staining, rinsing, and rapidly differentiated using 1% hydrochloric acid. The sections were then dehydrated in 80% ethanol, 90% ethanol, and absolute ethanol, washed with xylene and mounted with synthetic resin and examined under a light microscope (Olympus microscope, Japan). Histopathological evaluations were independently performed and reviewed under blinded conditions.

### Immunohistochemistry (IHC) analysis

4 µm tissue sections were fixed on slides by heating at 60°C, air-dried, and incubation at 60°C for 1 hour. For dewaxing xylene was used followed by rehydration in ethanol, and peroxidase activity was blocked with H₂O₂. After antigen retrieval, sections were incubated with 1% Bovine Serum Albumin-Transfer Buffer for 30 minutes, primary antibody (CD44, α-SMA and VIM) for 40 minutes, and anti-rabbit secondary antibody for 45 minutes (Mouse/Rabbit PolyVue Plus™ HRP/DAB Detection System, Diagnostic Biosystems). HRP (Horse reddish Peroxidase) substrate was added, followed by haematoxylin staining, dehydration, washing, and mounting with DPX (Dibutylphthalate Polystyrene Xylene) for microscopy (Olympus, Japan).

### Statistical analysis

Two-tailed unpaired t-tests were used to compare continuous variables between groups, assuming independence of samples, approximate normality, and equal variances. Analyses were performed in GraphPad Prism (RRID:SCR_002798), and results are reported as t-statistics, exact p-values, sample sizes, mean ± standard deviation (SD), and effect sizes with 95% confidence intervals. Survival outcomes were assessed using the Kaplan–Meier method, with group comparisons performed using the log-rank test in IBM SPSS Statistics v26 (RRID:SCR_002865). Analyses assumed independent survival times, non-informative censoring, and proportional hazards. Median survival with 95% confidence intervals and exact p-values are reported. Cohort-level variability and group separation were examined using Principal Component Analysis (PCA) implemented with the plot3D package in R (RRID:SCR_001905). Diagnostic performance of miR-1307-5p was evaluated using ROC curve analysis in GraphPad Prism, with the area under the curve (AUC) used as a measure of accuracy. Differential expression was visualised using volcano plots, where features with log₂ fold change ≥ 2 or ≤ –2 and adjusted p < 0.05 were considered significantly dysregulated. All experiments were performed in triplicate, and data are presented as mean ± SD.

## Results

### Identification of miR-1307-5p as a Clinically Relevant Predictor of Chemoresistance in OSCC

Salivary EVs were isolated and characterised by NTA, TEM, and flow cytometry. NTA indicated a concentration of 1.44 × 10^9^ particles/mL with a mean size of 36.9 ± 6.0 nm (Supplementary Fig. 1a). TEM analysis confirmed the spherical morphology substantiating a size range of 30–150 nm vesicles (Supplementary Fig. 1b). Flow cytometric analysis further validated the purity of the EV subpopulation by detecting the presence of tetraspanins (CD63, CD81, CD9) (Supplementary Fig. 1c).

Expression profile analysis demonstrated significantly elevated miR-1307-5p levels in chemo-non-responders (log₂FC: 3.34 ± 1.02; p < 0.001) compared to responders (log₂FC: 0.46 ± 1.6) (Fig. 1a). ROC analysis revealed significant predictive accuracy (AUC = 0.98; p < 0.001), while PCA plots distinctly separated non-responders, responders, and healthy controls, strengthening the discriminatory potential of salivary EV-derived miR-1307-5p (Fig. 1b,c). Further, to confirm these findings at the tissue level, in situ hybridisation (ISH) was performed on matched FFPE tumour sections from the same cohort. ISH revealed strong miR-1307-5p staining in chemo-non-responders, aligning with salivary EV expression (Fig. 1d,e).

**Figure 1:**
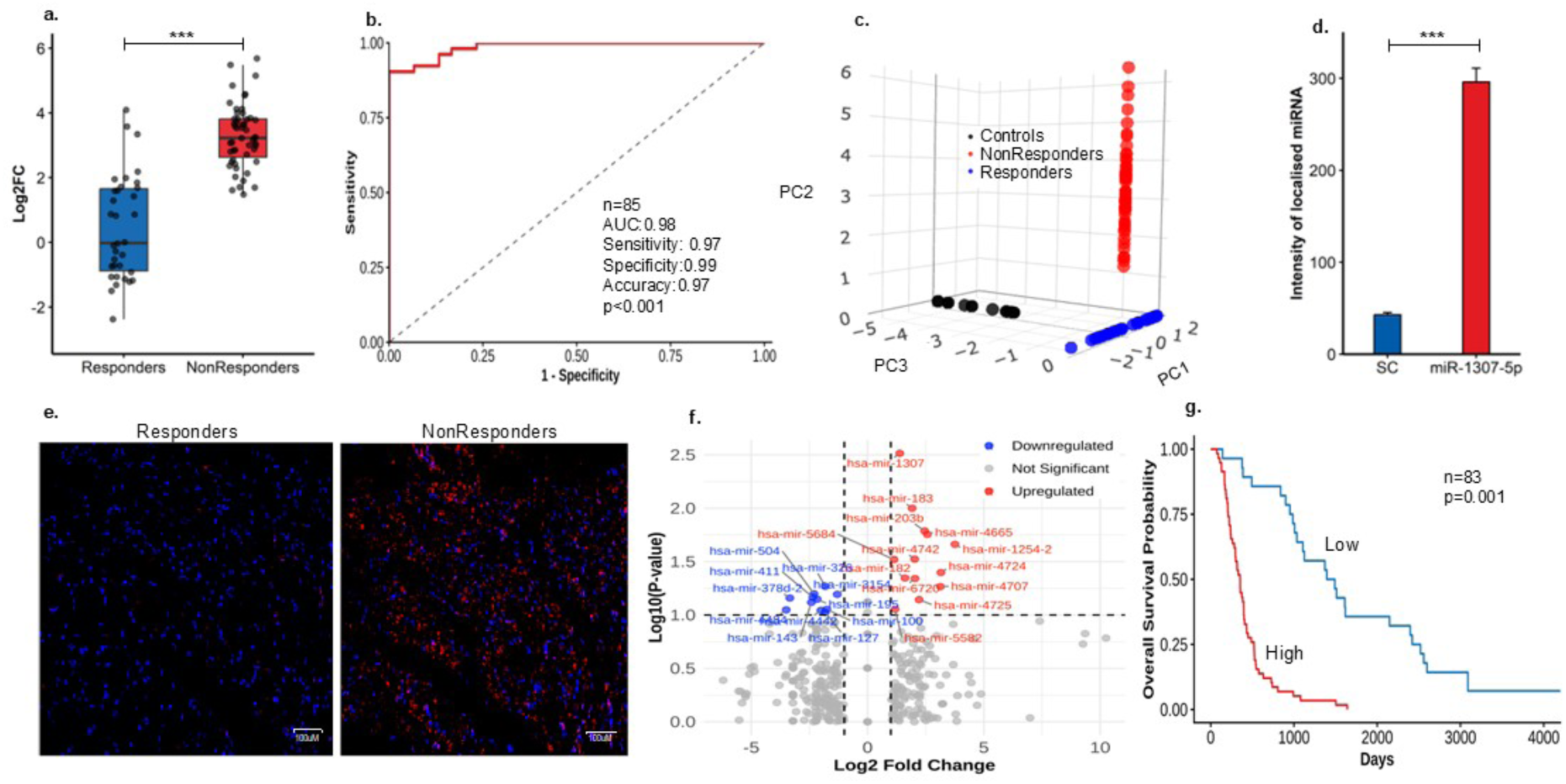
Salivary EV-Derived miR-1307-5p Predicts Chemotherapeutic Resistance in OSCC Patients. (a) Box-plot illustrating upregulated miR-1307-5p levels in salivary exosomes of chemo-non-responders (n=52) as compared to chemo-responders (n=33) using qPCR. Expression levels were normalised against U6 values, and relative quantification was performed using the ΔΔCt method. (b) ROC curve demonstrating the diagnostic accuracy of miR-1307-5p in OSCC patients (n=85), with AUC of 0.98 (p<0.01), 97% sensitivity, and 99% specificity. (c) PCA plot indicates distinct clustering between chemo-responders and non-responders, and controls (p<0.01) based on miR-1307-5p expression. (blue dots: chemo-responders, red dots: non-responders, black dots: controls) (d) Quantitative bar-graph representing miR-1307-5p and scramble-control (SC) expression in OSCC patient-derived tumour-sections assessed using ISH. (e) Subcellular localisation of miR-1307-5p in OSCC tumour tissues cells was determined by in situ hybridisation (ISH). Red staining (DIG-labelled probe miR-1307-5p), blue staining (DAPI). Scale bar: 100 µm (f) Volcano-plot depicting the differential expression of miRNAs in chemo-resistant OSCC patients from TCGA datasets. miR-1307-5p is significantly upregulated (p<0.001) among the top ten highly expressed miRNAs. The X-axis represents the log2FC, while the Y-axis shows the negative logarithm of the p-value (-log10 p-value). miRNAs with significant differential expression are highlighted (red), with threshold lines set at a p-value < 0.01 (indicated by the horizontal dashed line) and log2FC>2 (indicated by the vertical dashed lines) (g) Kaplan-Meier survival plot depicting upregulation of miR-1307-5p associated with poor survival outcomes of OSCC patients in TCGA_HNSC datasets (n=83). All experiments were conducted in triplicate, and error bars represent mean ± SD from three independent experiments. All statistical analyses in this figure are Student’s t-test unless specified. p<0.05 (*), p<0.01 (**), and p<0.001 (***). ROC: Receiver Operating Characteristic; AUC: Area under the curve; PCA: Principal Component Analysis.

To further validate miRNAs associated with chemoresistance, small RNA sequencing data profiling from TCGA–HNSC were analysed after segregating 83 OSCC patients who received complete therapy into chemo-responders and chemo-non-responders. The analysis identified 1,800 expressed miRNAs, of which 303 were differentially expressed between the two groups. Among these, 54 miRNAs were exclusively detected in chemo-non responders, applying the cut-offs for fold-change (FC: ± 2) and adjusted p-values (p< 0.05), 24 miRNAs remained significantly dysregulated (Fig. 1f). Notably, miR-1307-5p emerged as the most significantly upregulated marker in non-responder patients, highlighting its potential prognostic relevance. Consistent with our previous small RNA sequencing study (10), it was the only miRNA concordantly elevated in chemoresistant patients, underscoring its cross-platform robustness. Kaplan–Meier analysis further demonstrated significantly poorer outcomes in patients with high miR-1307-5p expression (p = 0.001), supporting its potential as a predictive biomarker of chemoresistance (Fig. 1g).

Collectively, this integrative clinical and translational analysis highlights the role of miR-1307-5p as a robust, reproducible, and non-invasive biomarker of chemoresistance, prompting to investigate its mechanistic and functional role in driving therapeutic refractoriness in OSCC.

### Targeting miR-1307-5p Reverses Malignant Traits in Cisplatin-Resistant CD44⁺ CSCs

Given the strong association of miR-1307-5p with chemo resistant patient cohorts, its functional and mechanistic relevance was investigated in CD44⁺ CSCs, an intrinsically resistant subpopulation in OSCC. Consistently, OECM1 cells showed significantly elevated CD44 surface expression (81.58 % ± 4.02, p=0.001) and transcript levels (log₂FC: 5.89 ± 0.17, p=0.001) compared with the negative control HEPG2 cells (Supplementary Fig. 2a,b). Further, CD44⁺ cells were treated with increasing concentrations of anti-miR-1307-5p (2.5–75nM) and scrambled control (SC). Anti-miR-1307-5p exhibited a dose-dependent cytotoxic effect with an IC₅₀ of 25nM, reducing cell viability to 53 ± 1.59% compared with SC (81 ± 2.43%; p < 0.01) (Fig. 2a). Further, qPCR results demonstrated a 30-fold reduction in the expression of miR-1307-5p levels at this dose (FC: 0.5 ± 0.5 vs. SC: 34 ± 1.7; p < 0.001) (Supplementary Fig. 2c). We analysed cell-cycle distribution, to determine the relevance of miR-1307-5p suppression on this chemo-resistant quiescent subpopulation. Anti-miR-1307-5p significantly increased the G2/M population (23.77 ± 1.36%) compared to SC (10.32 ± 0.48%; p < 0.001), indicating cell-cycle arrest and impaired proliferative capacity of the resistant CSCs (Fig. 2b; Supplementary Fig. 3a). Given the elevated sub-G0 population, we assessed apoptosis through annexin V-FITC/PI staining. The analysis showed marked increase in early (26.03 ± 1.35%) and late apoptosis (7.2 ± 0.38%) compared with SC (5.36 ± 0.2% and 2.76 ± 0.25%; p < 0.001) (Fig. 2c; Supplementary Fig. 3b). Morphological evaluation of PI-stained cells demonstrated apoptotic features of nuclear condensation and fragmentation (Supplementary Fig. 3c,d). JC-1 analysis further demonstrated mitochondrial membrane depolarisation (p < 0.001), confirming activation of intrinsic apoptosis (Fig. 2d; Supplementary Fig. 3e). Given that mitochondria are a major source of intracellular ROS and contribute to membrane depolarisation and intrinsic apoptotic signalling, we next assessed ROS levels. miR-1307-5p inhibition was associated with a marked increase in intracellular ROS in anti-miR-1307-5p–treated cells (9.89 ± 0.49, p = 0.0001) compared to scrambled controls (1.00 ± 0.05) (Fig. 2e). To assess whether miR-1307-5p regulates CSC aggressiveness, we evaluated the effect of its migratory and invasive potential. Anti-miR-1307-5p significantly reduced wound closure by 52% (2009 ± 76 µm vs. SC: 4144 ± 90 µm; p < 0.001) (Fig. 2f; Supplementary Fig. 3f), and invasion by 57% (43.3% ± 2.16% vs. SC: 10’0% ± 3%; p < 0.001) (Fig. 2h). In 3D spheroid assays, invasive protrusions were reduced by 21% (p < 0.001) (7.07 ± 1.00 vs. SC: 26.74 ± 1.3; p < 0.001) (Fig. 2g; Supplementary Fig. 3g). Given that CSCs modulate the tumour microenvironment via angiogenic signalling, we analysed endothelial tube formation using calcein AM staining. Anti-miR-1307-5p decreased tube network formation by 56% (142 ± 2.7 vs. SC: 252 ± 3.5; p < 0.001) (Fig. 2i; Supplementary Fig. 3h) and correspondingly reduced VEGF-α (FC: −5.1 ± 0.255) (Fig. 3c). Consistently, CAM assays demonstrated a marked reduction in vascular branching (15 ± 1.24 vs. 38 ± 2.62; p < 0.001) compared to the SC treated model system (Fig. 2j; Supplementary Fig. 3i). Collectively, these results indicate that miR-1307-5p sustains chemo resistant CSC phenotypes by maintaining mitochondrial integrity and redox homeostasis, whereas its inhibition disrupts this balance, leading to cell-cycle arrest, ROS-mediated apoptosis, and attenuation of tumour aggressiveness.

**Figure 2:**
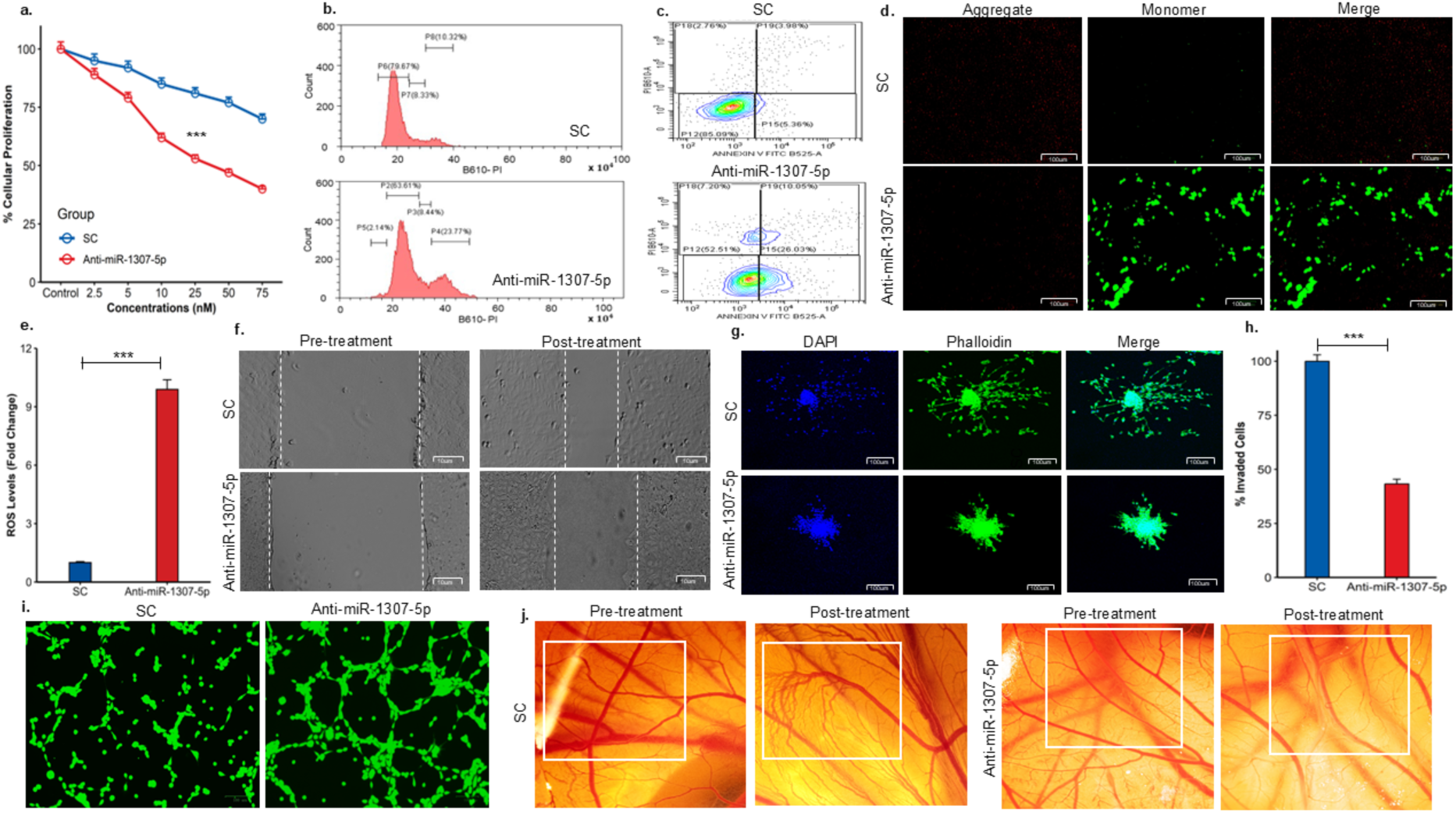
Anti-miR-1307-5p suppresses Proliferation, Induces cell-cycle arrest, Apoptosis, and Inhibits Invasion, Migration, Angiogenesis in CD44+ CSCs. (a) MTT assay showing reduced proliferation of CD44⁺ cells treated with anti-miR-1307-5p compared to scramble control (SC). The Y-axis represents percent proliferation normalised to untreated cells, and the X-axis shows concentrations (nM) of SC and anti-miR-1307-5p. (b) Flow-cytometry histograms of PI-stained CD44⁺ cells presenting DNA content distribution. PI fluorescence at 610 nm is plotted on the X-axis and cell count on the Y-axis. Cell-cycle quantification shows increased G₂/M arrest with a corresponding reduction in G₀/G₁ following anti-miR-1307-5p treatment. (c) Annexin-V/PI flow-cytometry scatter plots of CD44⁺ cells after transfection with SC or anti-miR-1307-5p. (d) Mitochondrial membrane potential (MMP) analysis (scale: 100 µm) displaying a shift from red (polarized mitochondria) to green (depolarized mitochondria), indicating MMP loss and mitochondrial dysfunction in anti-miR-1307-5p-treated cells. (e) Bar plot representing the quantification of ROS levels using DCFDA dye in anti-miR-1307-5p compared to SC. (f) Representative migration assay images (scale: 10 µm) showing reduced migratory capacity in anti-miR-1307-5p-treated cells (g) Confocal images of phalloidin- and DAPI-stained CD44⁺ spheroids (10×) demonstrating reduced invasive extensions following anti-miR-1307-5p treatment (h) Transwell invasion quantification showing decreased invasive ability of CD44⁺ CSCs treated with anti-miR-1307-5p compared to SC. (i) Calcein-AM fluorescence images of HUVEC tube formation in co-culture with treated CD44⁺ cells, showing diminished endothelial network development in the anti-miR-1307-5p group. (j) Ex-vivo CAM assay images showing reduced vascular branching and density after anti-miR-1307-5p treatment. Data represent mean ± SD (n=3). Statistical significance: p<0.05 (), p<0.01 (**), p<0.001 (***). SC: Scramble; PI: Propidium Iodide; MMP: Mitochondrial Membrane Potential; HUVECs: Human Large Vessel Endothelial Cells; CAM: Chick Chorioallantoic Membrane.

**Figure 3.**
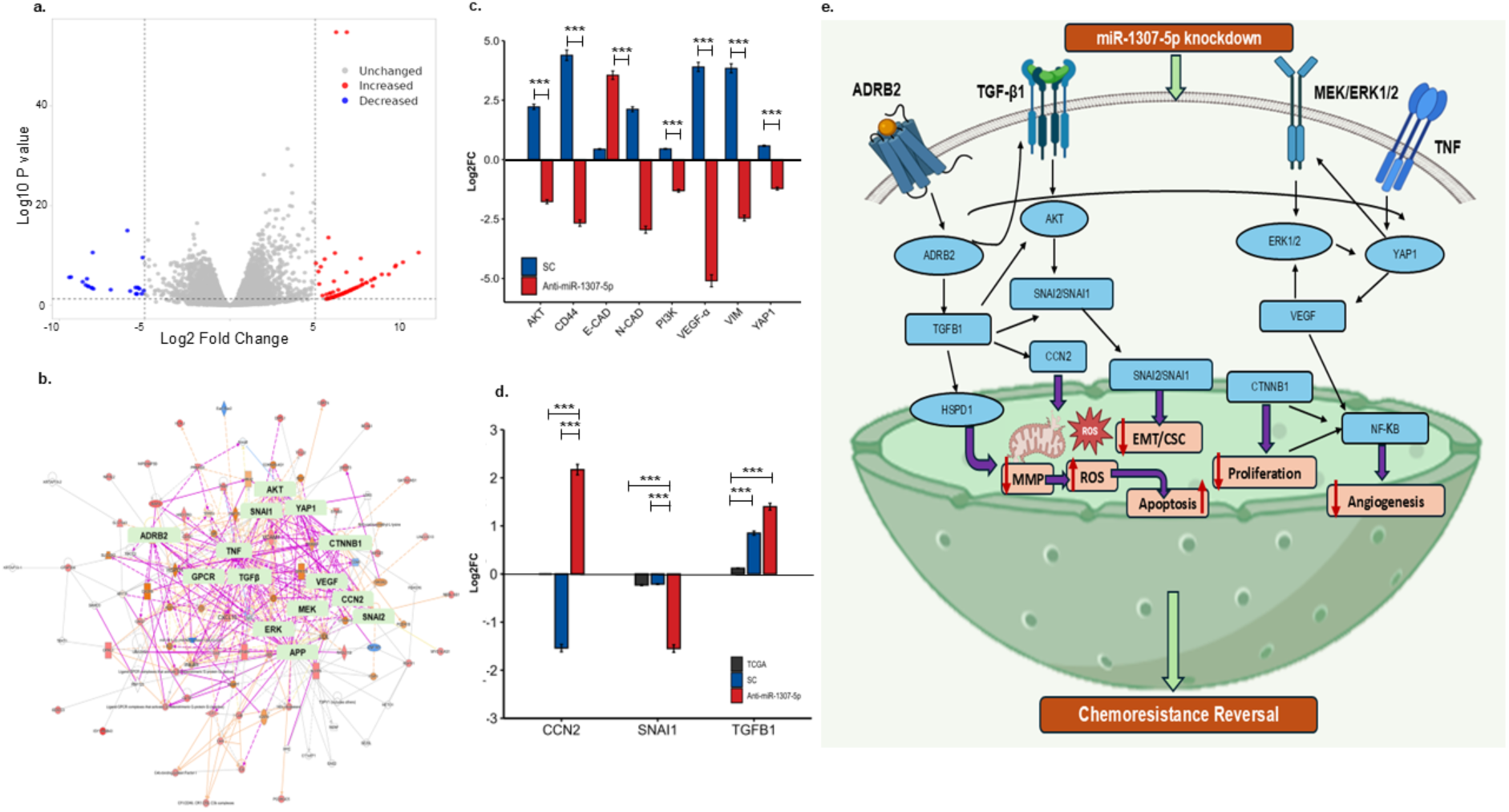
Integrated analysis of miR-1307-5p inhibition reveals transcriptional changes and pathway modulation associated with chemoresistance. (a) Volcano plot representing significant genes (Log2FC ± 2 and p-value<0.05) identified in anti-miR-1307-5p vs SC from a total of 16211 genes. Red dots indicate significantly upregulated genes, blue dots indicate significantly downregulated genes, and grey dots represent non-significant genes (based on thresholds). (b) Ingenuity Pathway Analysis (IPA) network of significantly altered genes, with key hub genes highlighted in green (bold) indicate their central role in regulatory signaling. (c) Expression levels (Log2FC) of genes regulating the proliferation, cell-cycle progression, apoptosis resistance, EMT and angiogenesis regulators including E-CAD, N-CAD, VIM, VEGF-A, CD44, AKT, YAP1 and PI3K following miR-1307-5p inhibition. Expression levels were normalised against β-actin values, and relative quantification was performed using the ΔΔCt method. (d) Bar plot illustrating the log2 fold change (Log2FC) of selected genes (CCN2, SNAI1, and TGFB1) in the SC and anti-miR-1307-5p-treated cohort, along with their concordance with the TCGA HNSCC dataset. (e) Schematic representation of the proposed molecular mechanism. miR-1307-5p knockdown modulates key signaling pathways including TGF-β, AKT, YAP1 and MEK/ERK, leading to regulation of EMT/CSC traits, proliferation, angiogenesis, and apoptosis. This ultimately contributes to reversal of chemoresistance. Error bars represent mean ± SD of three independent experiments. The statistical significance is indicated as: (*p<0.05; **p<0.01; ***p<0.001).

### Transcriptomic profiling reveals miR-1307-5p mediates chemoresistance via EMT and survival signalling pathways in OSCC

To delineate the molecular mediators underlying miR-1307-5p–driven chemoresistance, transcriptomic profiling was performed in anti-miR-1307-5p–treated CD44⁺ CSCs compared to SC (Fig. 3a). Using a read cutoff of >40, a total of 1,993 transcripts were uniquely detected in the treated population. Differential expression analysis (DESeq2) identified 109 genes that were significantly dysregulated compared to scrambled controls (|log₂FC| ≥ 5, adjusted p < 0.01). Network analysis of the 109 differentially expressed genes was performed using IPA. Genes were ranked by connectivity (node degree) based on experimentally validated and high-confidence predicted interactions, and hub genes were defined as those with >10 interactions within the reconstructed network (Fig. 3b). *TGFB1, TNF, APP, CTNNB1, YAP1, AKT, MEK, ERK, SNAI1, VEGF, ADRB2, SNAI2, CG, HSPD1, IgE,* and *CCN2* were identified as top hub genes from the network analysis. Subsequent PCR validation confirmed significant dysregulation of key candidates, including *VEGF* (log₂FC: −5.1 ± 0.255, p<0.001), *AKT* (log₂FC: −1.77 ± 0.088, p<0.001), *YAP1* (log₂FC: −1.22 ± 0.061, p<0.001), and *PI3K* (log₂FC: −1.31 ± 0.065, p<0.001), in anti-miR-1307-5p–treated CD44⁺ CSCs compared to SC (Fig. 3c). Cross-referencing with TCGA OSCC datasets further revealed that *SNAI1 (*log₂FC: *-1.55* ± 0.07, p<0.001), *CCN2 (*log₂FC: *2.17* ± 0.10, p<0.001), and *TGFB1 (*log₂FC: *1.4* ± 0.07, p<0.001) displayed consistent expression patterns in patient cohorts and inverse regulation upon miR-1307-5p inhibition suggesting suppression of EMT-associated programs (Fig. 3d). Consistent with this, EMT reversal was confirmed at the transcript level by increased E-cadherin expression (log₂FC: 1.5 ± 0.07) and reduced N-cadherin (log₂FC: −1.61 ± 0.08) and vimentin levels (log₂FC: −2.46 ± 0.123) (Fig. 3c).

Collectively, these findings indicate convergence on major oncogenic signaling axes, including PI3K–AKT, MAPK/ERK, and YAP-associated pathways, which govern proliferation, cell-cycle progression, apoptosis resistance, EMT, and angiogenesis (Fig. 3e). Moreover, these results align with the observed functional reprogramming following miR-1307-5p inhibition, underscoring its role as a central regulator of CSC-driven chemoresistance networks.

### Targeting miR-1307-5p Restores Low-Dose Cisplatin Efficacy and Restrains CD44⁺ CSC–Driven Tumour Growth In Xenograft Model

To determine whether miR-1307-5p inhibition could reverse chemoresistance, we evaluated the effects of anti-miR-1307-5p on cisplatin response in CD44⁺ subpopulation. CD44⁺ cells exhibited a significant reduction in viability (52.9 ± 1.58%; p < 0.01) when co-treated with 0.1µM cisplatin and 10nM anti-miR-1307-5p, compared with SC treated cells (68 ± 2.05%; p < 0.01), indicating a ∼25-fold decrease in the cisplatin dose required to induce cytotoxicity in this chemo resistant population (Fig. 4a). Moreover, qPCR analysis confirmed substantial post-treatment suppression of miR-1307-5p (FC: 0.19; p < 0.01), indicating re-sensitisation attributed to miRNA inhibition rather than nonspecific cytotoxicity (Supplementary Fig. 4a). To understand the implications of enhanced cisplatin sensitivity, we assessed its effect on cell-cycle progression and apoptotic activation. Anti-miR-1307-5p along with low dose cisplatin induced a substantial G2/M arrest (31.44 ± 0.94%) compared to SC + cisplatin treated cells (12.45 ± 0.37%; p < 0.001), suggesting impaired mitotic progression (Fig. 4b; Supplementary Fig. 4b). These findings align with cisplatin’s mechanism of action, as its DNA crosslinking effect predominantly impedes cells at the G2/M checkpoint. Further, a sub-G0/G1 peak prompted apoptosis analyses, where PI staining revealed characteristic apoptotic morphological alterations (Fig. 4d; Supplementary Fig. 4c) and JC-1 assays demonstrated a significant reduction in mitochondrial membrane potential (red/green ratio: 0.5 ± 0.004 vs. 3.21 ± 0.09; p < 0.001), confirming intrinsic apoptotic activation (Fig. 4c; Supplementary Fig. 4d). Further miR-1307-5p inhibition was also associated with a marked increase in ROS levels in anti-miR-1307-5p+Cis treated cells (14.51 ± 0.72, p=0.0001) compared to SC+Cis (2.9 ± 0.14) (Fig. 4e).

**Figure 4:**
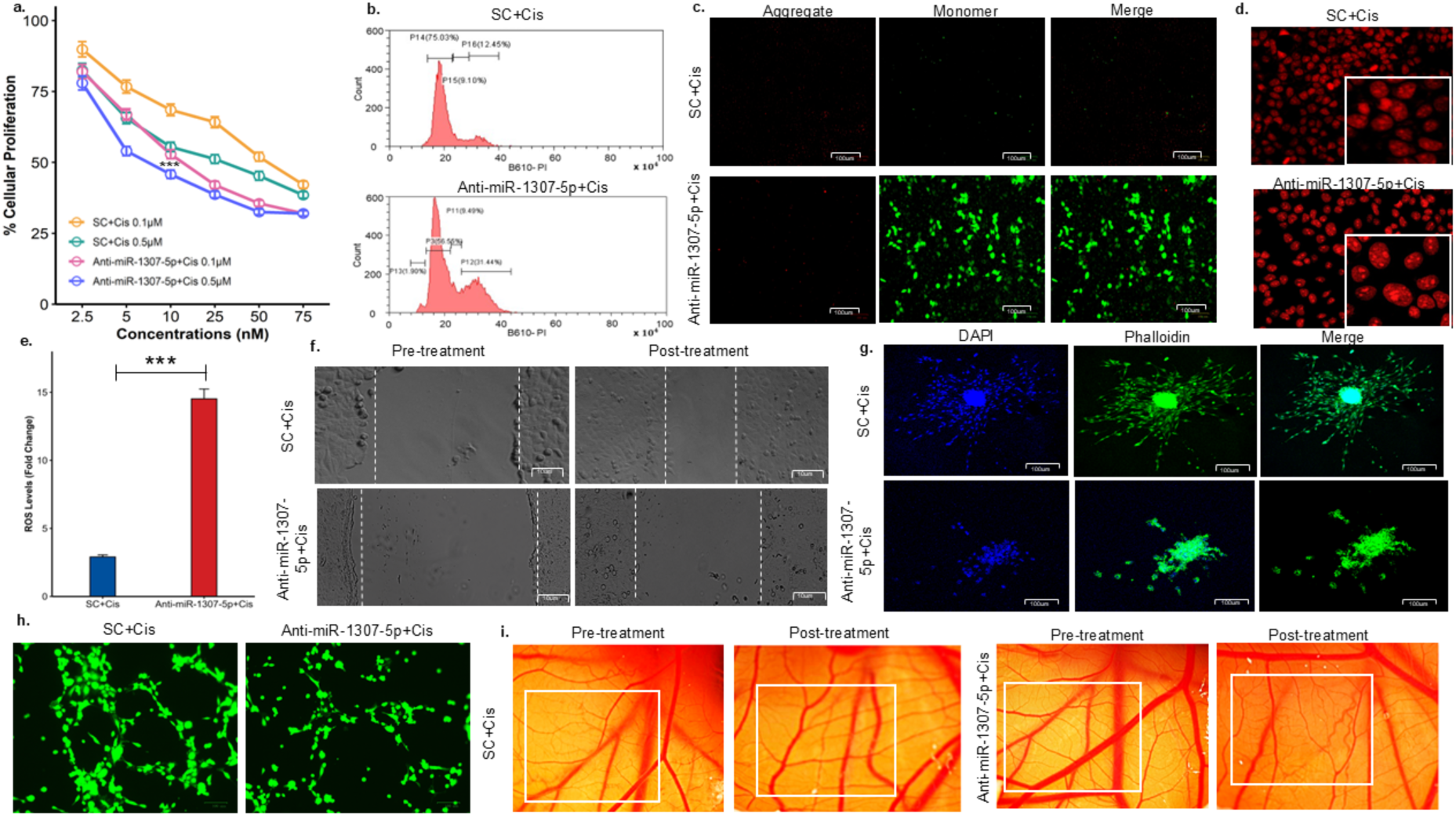
miR-1307-5p Suppression Restores Cisplatin Sensitivity in CD44+ CSCs. (a) MTT assay demonstrated reduction in the proliferation of CD44⁺ cells treated with anti-miR-1307-5p+Cis compared with SC+Cis, with percentage proliferation normalised to untreated controls across increasing concentrations. (b) Flow-cytometry analysis of PI-stained cells revealed that anti-miR-1307-5p+Cis induced a notable shift in DNA content, shown by a significant increase in the G2/M population accompanied by a decrease in G0/G1 relative to SC+Cis. (c) Mitochondrial membrane potential assessment (scale: 100 µm) demonstrated a transition from red, indicating polarised mitochondria, to green, indicating depolarised mitochondria, in anti-miR-1307-5p+Cis, reflecting mitochondrial dysfunction and apoptotic activation. (d) Fluorescence imaging of PI-stained CD44⁺ cells (scale: 35 µm) showed more pronounced apoptotic morphology, including nuclear condensation and fragmentation, under anti-miR-1307-5p+Cis compared with SC+Cis. (e) Bar plot representing the quantification of ROS levels using DCFDA dye in anti-miR-1307-5p+Cis compared to SC+Cis. (f) Wound-healing assay images (scale: 10 µm) demonstrated reduced migratory ability in anti-miR-1307-5p+Cis-treated cells (g) Confocal images of phalloidin- and DAPI-stained CD44⁺ spheroids at 10× magnification displayed diminished invasive protrusions with anti-miR-1307-5p+Cis, and quantitative analysis verified substantial suppression of spheroid invasiveness compared with SC+Cis. (h) Fluorescence images of calcein-AM-stained tube formation in HUVECs co-cultured with CD44⁺ cells illustrated significantly reduced endothelial network formation after anti-miR-1307-5p+Cis treatment. (i) *Ex-vivo* CAM assay images showed decreased vascular branching and density in anti-miR-1307-5p+Cis-treated samples relative to SC+Cis. Data are presented as mean ± SD from three independent experiments. Statistical significance: p<0.05 (*), p<0.01 (**), p<0.001 (***). SC: Scramble; Cis: Cisplatin; PI: Propidium Iodide; MMP: Mitochondrial Membrane Potential; HUVECs: Human Large Vessel Endothelial Cells; CAM: Chick Chorioallantoic Membrane Model.

Notably, even though anti-miR-1307-5p standalone could also induce apoptosis and G2/M arrest, the addition of low dose cisplatin along with reduced concentration of anti-miR-1307 resulted in a significantly amplified outcome. These findings indicated that the synergistic effect of this combination could reprogramme resistant CSCs into a therapy-responsive state. Given that CSC-mediated chemoresistance is closely associated with invasion and EMT, we assessed the effect of combination therapy on aggressiveness. Co-treatment significantly inhibited migration (497 ± 14.91 µm vs. SC + cisplatin: 3141 ± 94.23 µm; p < 0.01) (Fig. 4f; Supplementary Fig. 4e), reduced invasion by 50% (32% ± 0.97% vs. 82% ± 2.48%; p < 0.001) (Supplementary Fig. 4f), and decreased 3D spheroid invasiveness (6.67 ± 0.36 vs. 26 ± 2%; p < 0.001) (Fig. 4g; Supplementary Fig. 4g). These effects were substantially greater than anti-miR-1307-5p alone, demonstrating enhanced suppression of CSC-driven metastatic traits. Further co-treatment reduced endothelial tube formation by 40% (81 ± 3.4 vs. 200 ± 3.5; p < 0.001) (Fig. 4h; Supplementary Fig. 4h) and decreased vascular branching in CAM assays by 29% (10 ± 0.5 vs. 35 ± 1.25; p < 0.001) (Fig. 4i; Supplementary Fig. 4i), exceeding the anti-angiogenic effects of miR-1307-5p inhibition alone.

Owing to the robust chemo-sensitisation observed *in vitro* and *ex vivo*, we evaluated the therapeutic impact of miR-1307-5p suppression in an orthotopic OSCC mouse model. CD44⁺ CSCs transduced with sh-miR-1307-5p and sh-control lentiviral particles were orthotopically injected into the buccal mucosa of NOD-SCID mice, followed by subsequent treatment with low-dose cisplatin (2 mg/kg). Tumour volumes were markedly reduced in the sh-miR-1307-5p + cisplatin group (4 ± 0.12 mm³; p < 0.0001) compared with sh-control + cisplatin (7 ± 0.21 mm³), sh-miR-1307-5p standalone (6 ± 0.18 mm³), and untreated controls cohorts (9 ± 0.27 mm³) (Fig. 5a,b). Notably, neither cisplatin nor sh-miR-1307-5p alone produced significant tumour reduction, emphasising the significant effect of combinatorial treatment in inducing tumour regression by targeting chemo-resistant population and restoring cisplatin sensitivity *in-vivo.* Furthermore, histopathological and immunohistochemical analyses revealed enhanced tumour cell death, partial restoration of differentiated epithelial morphology, and a marked reduction in CD44, ɑ-SMA and VIM expression, particularly in the combination group (Fig. 5c,d).

**Figure 5:**
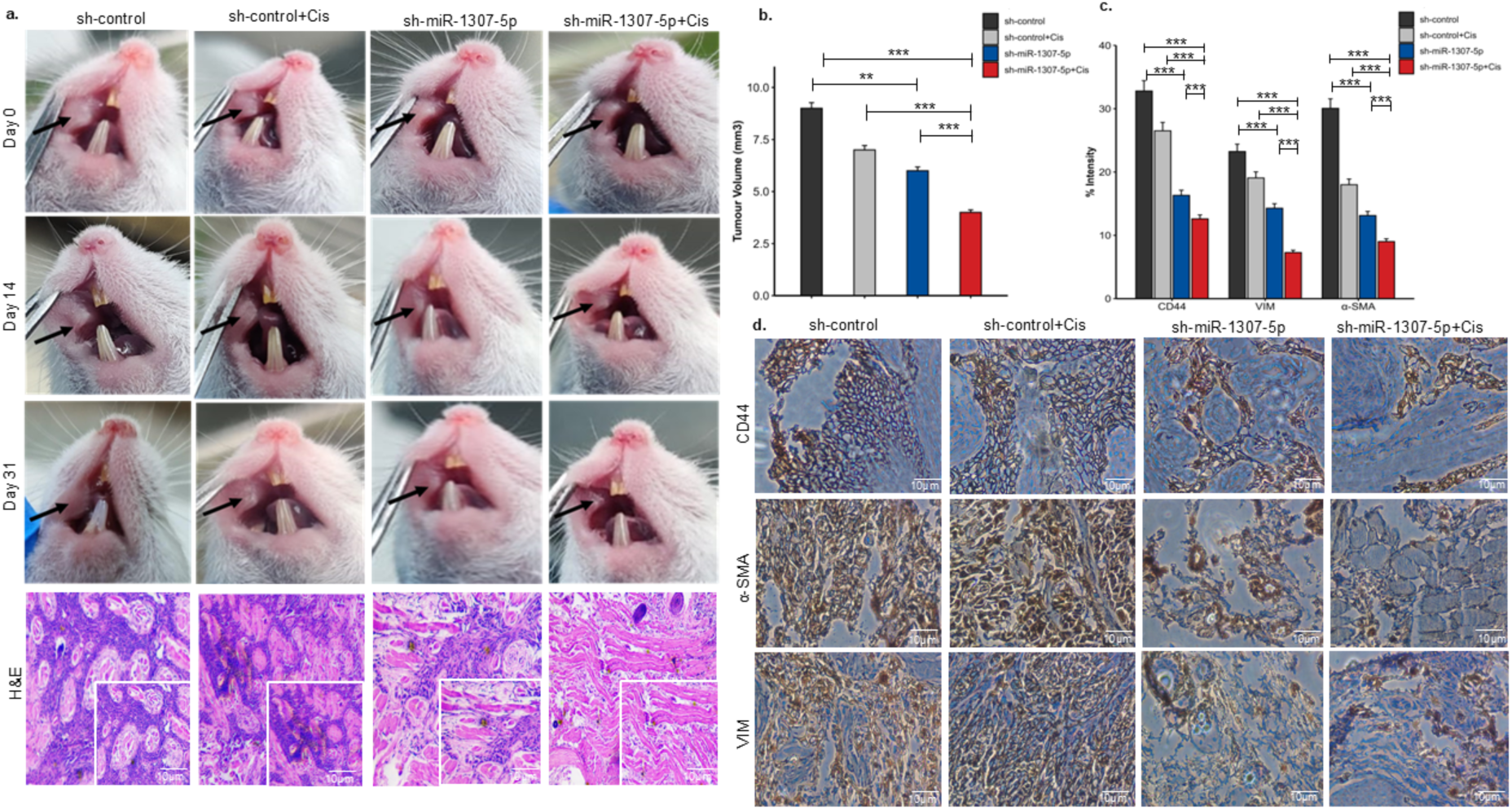
Inhibiting miR-1307-5p Decreases Tumour Growth and Enhances Cisplatin Efficacy in a Xenograft Mouse Model. (a) Representative images of the tumour morphology following treatment with sh-control, sh-control+Cis, sh-miR-1307-5p, and sh-miR-1307-5p+Cis. Tumours in the sh-miR-1307-5p and sh-miR-1307-5p+Cis-treated mice exhibit a visible reduction in size compared to SC and SC+Cis, respectively. H&E-stained tumour sections reveal decreased cell density, nuclear alterations, and disrupted tumour architecture, indicating reduced malignancy in the tumors of sh-miR-1307-5p and sh-miR-1307-5p+Cis as compared to sh-control, sh-control+Cis. (b) Tumour volume measurements demonstrate a significant reduction in sh-miR-1307 and sh-miR-1307+Cis groups compared to sh-control and sh-control+Cis. (c) Quantification of IHC staining for CD44, α-SMA and VIM expression (percentage of positive stained marker area) shows a marked decrease of CD44, α-SMA and VIM expression in treated groups, indicating suppression of CD44+ cancer stem cell populations. (d) IHC staining for CD44, α-SMA and VIM (brown) demonstrates a significant reduction in the expression in sh-miR-1307-5p and sh-miR-1307-5p+Cis-treated tumours. Error bars represent mean ± SD of three independent experiments. The statistical significance is indicated as: (*p<0.05; **p<0.01; ***p<0.001). sh-control-Scramble-control, Cis-Cisplatin.

Thus, suppression of miR-1307-5p disrupts resistant traits, while combination with low-dose cisplatin forces CSCs into a therapy-responsive state. This synergy restores cisplatin efficacy at reduced doses, positioning miR-1307-5p as a promising therapeutic target to overcome chemoresistance in OSCC.

### Exosomal Transfer of Anti-miR-1307-5p Reprogrammes Resistant CSCs Towards a Therapy-Sensitive State

To determine whether EVs could adoptively transfer the chemo-sensitising to a chemoresistant population, we assessed whether EVs derived from anti-miR-1307-5p–treated CD44⁺ CSCs could reprogram untreated resistant CSCs toward a chemo-sensitive phenotype. Confocal imaging of labelled EVs confirmed efficient uptake by CD44⁺ CSCs, and qPCR analysis demonstrated a marked reduction of miR-1307-5p within EVs derived from anti-miR-treated cells (FC: 0.25 ± 0.01 vs. SC EVs: 2.82 ± 0.20; p < 0.02) (Supplementary Fig. 6a). Further, resistant CSCs were subsequently co-cultured with these EVs to assess functional impact. Dose-response assays identified an IC₅₀ of 7.7 × 10⁶ particles/mL, associated with a 45% reduction in cell proliferation of CD44+ CSCs compared to vehicular control derived EVs (48.11 ± 1.45 vs. SC-EVs: 87.95 ± 2.63; p < 0.01) (Fig. 6a). EVs derived from anti-miR-1307-5p treated cells induced significant G2/M arrest (22.53 ± 0.41 vs. SC-EVs: 13.77 ± 0.67; p < 0.001) (Fig. 6b; Supplementary Fig. 6b). Further, they demonstrated marked increase in early apoptosis (Annexin V⁺/PI⁻: 9.56 ± 0.28 vs. 2.42 ± 0.07; p < 0.0001), along with morphological changes on PI staining—including chromatin fragmentation and nuclear condensation (Fig. 6c; Supplementary Fig. 6c,d). These findings confirm that EV-mediated miR-1307-5p suppression reproduces the mirroring of direct anti-miRNA effects on cell cycle and intrinsic apoptotic responses.

**Figure 6:**
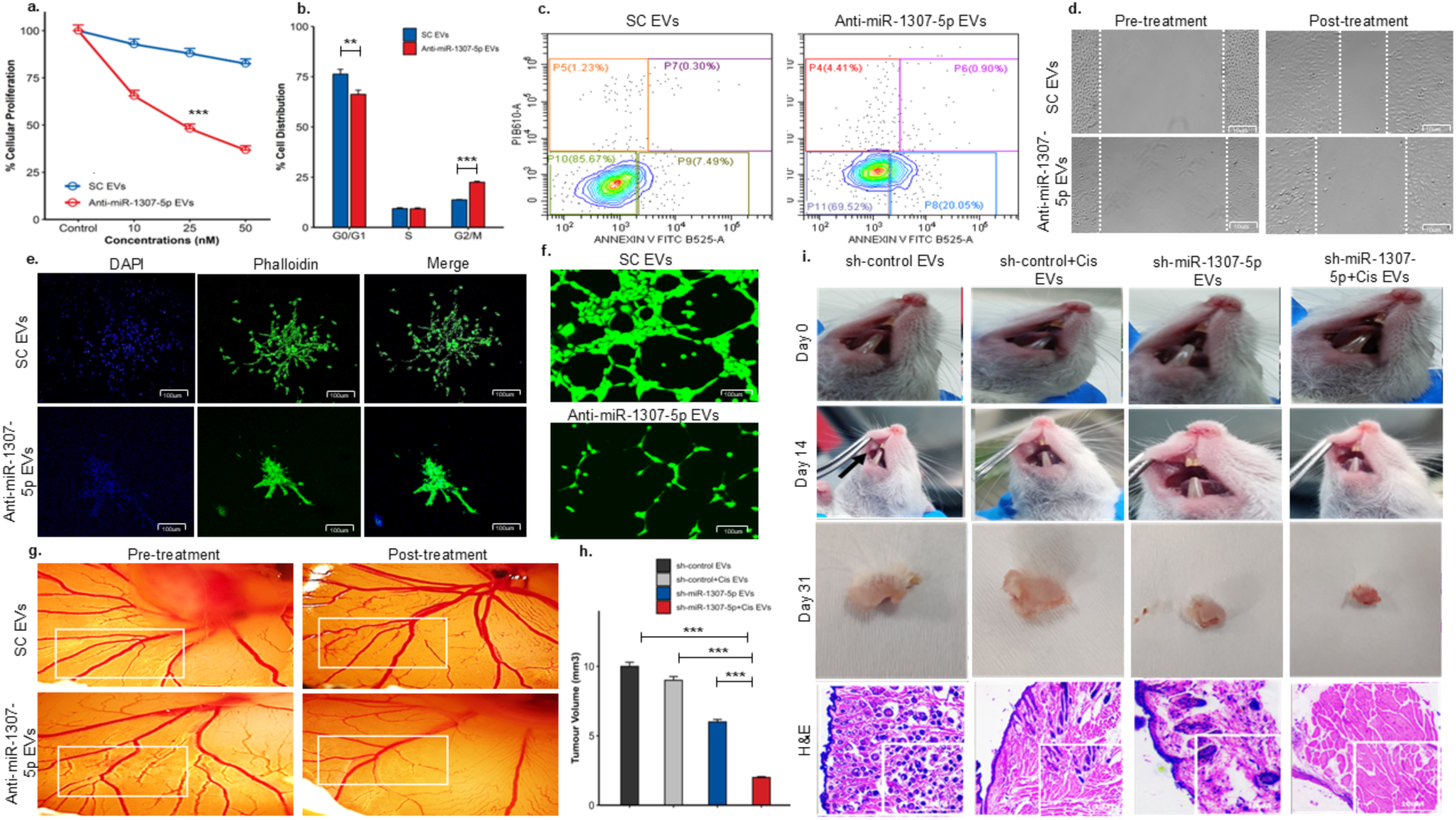
EV-mediated Delivery of anti-miR-1307-5p Inhibits Tumour Progression and Improves the Chemosensitivity in CSCs. (a) MTT assay showing reduced proliferation of CD44+ cells following treatment with anti-miR-1307-5p EVs. Y-axis: %proliferation of CD44+ cells, X-axis: concentration (nM). (b) Bar graph quantifying flow cytometry results, demonstrating a significant change in G0/G1 and G2/M phases in anti-miR-1307-5p EVs treated cells as compared to SC EVs (c) Flow cytometry scatter plot for Annexin-V/PI stained CD44+ cells after co-culture with anti-miR-1307-5p EVs as compared to SC EVs with anti-miR-1307-5p EVs showing a significant increase of early apoptotic population (Annexin V+/PI−) for treated cells (d) Microscopic images showing the decrease in migratory ability of CD44+ cells when treated with anti-miR-1307 EVs as compared to SC EVs (scale: 10µM) (e) Confocal images illustrating reduction in the invasive potential of CD44+ spheroids treated with anti-miR-1307-5p EVs compared to SC EVs. Spheroids were stained with phalloidin (green) and DAPI (blue); imaged at 10X magnification. (f) Representative images showing reduced angiogenic potential of HUVECs when co-cultured with anti-miR-1307-5p EVs as compared to SC EVs (g) CAM Images showing disrupted vascular development after anti-miR-1307 EVs treatment. (h) Bar-graph depicting reduction in tumour volume for sh-miR-1307 EVs+Cis treated mice as compared to SC EVs+Cis. (i) NOD-SCID xenograft mice model depicting a decrease in the tumour size when injected with sh-miR-1307 EVs+Cis in the buccal cavity region as compared to SC EVs+Cis. H&E staining of tumour sections shows reduced tumour cell density, altered nuclear morphology, and increased necrosis in sh-miR-1307 EVs+Cis treated mice. The data was normalised to controls, experiments were performed in triplicate and error bars represent mean ± SD. The statistical significance is indicated as: (*p<0.05; **p<0.01; ***p<0.001). HUVECs: Human Large Vessel Endothelial Cells, CAM: chick chorioallantoic membrane model, sh-control- Scramble-control, Cis- Cisplatin.

anti-miR-1307-5p treated EVs also impaired CSC aggressiveness with a 45% decrease in migratory potential (66.5 ± 2.02 µm vs. 94.25 ± 2.82 µm; p < 0.001), 88% reduction in invasive capability (11.9 ± 0.36 vs. 92 ± 2.76; p < 0.001), and 30% decline in the invasive index of 3D spheroids matrix (7.97 ± 0.23 vs. 18.68 ± 0.55; p < 0.01) (Fig. 6d,e; Supplementary Fig. 6e,f,g). EV-mediated inhibition also had a significant impact on the angiogenic potential of these cells by reducing endothelial tube formation by 43% (38 ± 1.14 vs. 67 ± 2.01; p < 0.001) and CAM vascular branching by 52% (20 ± 1.24 vs. 42 ± 1.62; p < 0.001) (Fig. 6f,g; Supplementary Fig. 6h,i).

To evaluate therapeutic potential of EVs *in-vivo*, NOD-SCID mice were orthotopically injected with EVs derived from anti-miR-1307-5p–treated and scrambled control CSCs. Tumour volumes were significantly reduced (6 ± 0.18 mm³ vs. SC-EVs: 10 ± 0.3 mm³; p < 0.001) (Fig. 6h). Histopathology revealed reduced tumour cell density, altered nuclear morphology, and enhanced necrosis. When combined with low-dose cisplatin (2 mg/kg), anti-miR-1307-5p treated EVs amplified tumour regression (2 ± 0.06 mm³ vs. SC-EVs + cisplatin: 9 ± 0.27 mm³; p < 0.001). H&E staining showed a striking shift toward differentiated, non-malignant morphology (Fig. 6i), suggesting reversal of CSC-driven tumorigenesis *in vivo*.

These findings show that EV-mediated delivery of anti-miR-1307-5p effectively recapitulates cellular reprogramming, suppresses CSC hallmarks, and, in combination with cisplatin, induces strong tumour regression. The close concordance with direct anti-miR silencing confirms the role of miR-1307-5p–enriched EVs in mediating chemoresistance and highlights EV-based anti-miR-1307-5p delivery as a promising strategy to overcome resistance in OSCC.

## Discussion

Chemoresistance poses a significant challenge in the management of OSCC, attributed to CD44⁺ CSC subpopulation, which possess enhanced survival mechanisms and contribute to tumour relapse after administering conventional therapeutic regimen. In this study, we identify the crucial role for salivary EV–derived miR-1307-5p in mediating chemoresistance, demonstrating both its clinical relevance as a biomarker and its mechanistic contribution to CSC-driven therapeutic failure. Combining patient salivary EV profiling, and TCGA analysis, we demonstrated that miR-1307-5p is consistently upregulated in chemoresistant patients and clinically associated with poor overall survival. These findings position miR-1307-5p as a robust predictor of chemoresistance and a promising tool for non-invasive treatment stratification. Studies identified miR-1307-5p as an oncogenic miRNA across multiple cancers, including lung, ovarian and liver malignancies (17-19). Notably, it is strongly upregulated in chemoresistant ovarian cancer cells and directly drives resistance (20). However, its role in OSCC chemoresistance, especially its mechanistic function and EV-associated involvement has never been explored. Our work is the first to demonstrate and mechanistically define miR-1307-5p, both cellular and EV-borne, as a driver of chemoresistance in OSCC, underscoring its clinical relevance and therapeutic promise.

Mechanistically, inhibition of miR-1307-5p reversed multiple CSC hallmarks including quiescence, apoptotic resistance, invasiveness, EMT, angiogenesis and restoring cisplatin sensitivity across *in-vitro*, *ex-vivo*, and *in-vivo* systems. These findings align with the prior observations in lung adenocarcinoma, hepatocellular carcinoma, osteosarcoma, and breast cancer (19, 21-23), where miR-1307 promoted aggressive phenotypes. Importantly, miR-1307-5p inhibition sensitised CSCs to low-dose cisplatin, yielding a stronger therapeutic response than standard inhibitory concentration of cisplatin, which could have high translational relevance given platinum-related toxicity. These results align with broader evidence establishing miRNAs as critical regulators of drug response: miR-148b enhances cisplatin sensitivity (23), multiple miRNAs modulate 5-FU and oxaliplatin resistance in colorectal cancer (6, 24-32), and EMT-associated miRNAs such as miR-221, miR-93, and miR-27a promote resistance across cancers (33-35). Collectively, these studies highlight the inevitable role of miRNAs in governing therapeutic outcomes. However, these studies focus on bulk tumour cells and rely on high clinical drug doses. In contrast, our study reveals miRNA-driven chemoresistance and its role in targeting CSC subpopulation under dose-sensitive, clinically relevant treatment conditions addressing a critical unmet gap.

To delineate the mechanistic basis of miR-1307-5p–mediated chemoresistance, transcriptomic profiling followed by IPA identified a convergent network of oncogenic hubs, integrating EMT, CSC maintenance, and mitochondrial homeostasis. The TGFB1–SNAI2–CTNNB1 axis is a well-established driver of EMT and CSC plasticity in lung and head and neck cancers, promoting invasion, stemness, and therapeutic resistance (36-38). Concurrently, AKT and MEK/ERK signaling pathways sustain CSC survival and self-renewal, while TNF–NF-κB signaling reinforces inflammatory programs that contribute to aggressive, drug-resistant phenotypes in head and neck cancers (39-41). YAP1 has emerged as a critical regulator of EMT and CSC traits in OSCC, integrating mechanical and metabolic cues to support tumour progression and therapy resistance (42, 43). In parallel, mitochondrial regulators such as HSPD1 maintain cellular bioenergetics and protect against stress-induced apoptosis, thereby preserving CSC fitness under chemotherapeutic stress (44). Silencing miR-1307-5p disrupted these interconnected pathways, leading to suppression of EMT and CSC-associated transcriptional programs, accompanied by mitochondrial destabilisation, increased ROS accumulation, and loss of mitochondrial membrane potential (45-47). Elevated ROS levels and mitochondrial dysfunction are known to impair CSC maintenance and sensitise OSCC cells to therapy (48, 49). Moreover, attenuation of YAP1- and AKT-driven survival signaling further lowers the threshold for CSC survival under stress conditions (50, 51). Collectively, these findings position miR-1307-5p as a central regulator of EMT–CSC–mitochondrial crosstalk in OSCC, where its inhibition disrupts redox balance and stemness-associated survival pathways, ultimately reprogramming resistant CSCs into a therapy-responsive state and reversing chemoresistance.

A novel and significant aspect of this study is the demonstration that EVs can significantly transmit therapy-sensitive phenotype. EVs from anti-miR-1307-5p–treated CSCs recapitulated the overall chemo-sensitising phenotype induced by direct knockdown in both *in-vitro* and *in-vivo* models. The *in-vivo models* demonstrated reduced CD44 levels and renewed cisplatin responsiveness, validating that miRNAs packaged in EVs have the capability to transmit these signals to neighbouring cells. This supports the concept that tumour-derived EVs function as mediators of resistance or sensitivity within the tumour microenvironment. Studies of exosomal miRNAs promoting resistance in HNSCC such as miR-21-5p, miR-7704, and miR-3960 by modulating various signalling pathways have been reported. Therapeutic exosomes carrying miR-155 inhibitors or ovatodiolide have similarly restored cisplatin sensitivity in preclinical models. Thus, EV-encapsulated anti-miRNA therapeutics offer a clinically translatable strategy for targeted and stable nucleic acid delivery, enabling effective modulation of drug resistance pathways while reducing off-target effects and systemic toxicity. This study provides the first evidence that EV-mediated delivery of anti-miR-1307-5p can induce tumour regression *in-vivo*, representing a translationally viable therapeutic modality. Given the stability and accessibility of salivary EVs, the clinical translation of miR-1307-5p as both a biomarker and therapeutic target presents a compelling opportunity for minimally invasive monitoring and precision therapy in OSCC.

Although transcriptomic analyses identified key regulatory hubs, these findings remain largely associative, and comprehensive validation of direct miR-1307-5p targets is required. Additionally, EV heterogeneity and challenges in large-scale production and standardisation may impact clinical translation. Nevertheless, EV-mediated delivery enables horizontal transfer of therapy-sensitive phenotypes, offering a unique strategy to reprogram resistant tumour ecosystems.

## Conclusion

In summary, our findings position EV-derived miR-1307-5p as a pivotal regulator and clinically relevant biomarker of chemoresistance in OSCC. Functional inhibition of miR-1307-5p either directly or via EV-mediated strategies effectively restores cisplatin sensitivity and abrogates aggressive CSC phenotypes. These results not only underscore its central role in driving therapeutic resistance but also establish miR-1307-5p as a compelling and actionable target for overcoming drug resistance in OSCC.

## List of Abbreviations

OSCC: Oral Squamous Cell Carcinoma
EVs: Extracellular Vesicles
miRNAs: micro-RNAs
CSCs: Cancer Stem Cells
CD44: Cluster of Differentiation 44
TCGA: The Cancer Genome Atlas
ISH: *in-situ* hybridisation
NTA: Nanoparticle-Tracking Analysis
HUVEC: Human Umbilical Vein Endothelial Cells
CAM: Chorioallantoic Membrane
IHC: Immunohistochemistry
H&E: Haematoxylin and Eosin
PFA: Paraformaldehyde
NBF: Neutral Buffered Formalin
HRP: Horseradish Peroxidase
DPX: Dibutyl phthalate Polystyrene Xylene
ROC: Receiver Operating Curve
PCA: Principal Component Analysis
EMT: Epithelial-to-Mesenchymal Transition

## Declarations

### Ethics approval and consent to participate

All procedures of human samples were conducted after the approval of the HCG Aastha Cancer Centre’s Ethics Committee (ECR/92lInst/GJ/2013/RR-19) for human subject research. This study also complied with the guidelines set forth by the Declaration of Helsinki (2008). All patients provided written informed consent for their participation in the study and their identities have been anonymized. The in-vivo study was approved by the Animal Ethics Committee of IISER, Pune (IISER-PUNE/IAEC/2023_03/05).

### Consent for Publication

All the authors have read and approved the final manuscript for publication.

### Availability of Data and Materials

The data generated in this study are available within the article and its supplementary data files.

### Author contributions

A.P.- Data curation, Formal analysis, Methodology, Validation, Visualisation, Writing – original draft. V.P.- Data curation, Formal analysis, Methodology, Validation, Visualisation. S.L.- Data curation, Formal analysis, Methodology, Validation. K.P.- Conceptualisation, Supervision. D.M.- Supervision. J.T.- Methodology. P.S.- Validation. B.P.- Methodology. K.J.- Visualisation. D.B.-Visualisation. V.T.- Conceptualisation, Investigation, Supervision, Writing – review & editing. S.P.- Conceptualisation, Investigation, Supervision, Resources, Funding acquisition, Writing –original draft.

## Acknowledgments

This work was partially supported by DBT/Wellcome Trust India Alliance Fellowship/Grant [grant number IA/E/20/1/505666] awarded to SP. The authors acknowledge DBT Sahaj for providing the resources necessary to carry out the flow cytometric analysis. The DBT/Wellcome Trust India Alliance Fellowship/Grant [grant number IA/E/20/1/505666] awarded to SP is highly acknowledged. AP acknowledges the Council of Scientific and Industrial Research (CSIR), Government of India, for the award of a Senior Research Fellowship. Lady Tata Memorial Trust (LTMT) is acknowledged for supporting SL with a Junior Research Fellowship.

We thank HCG Aastha Cancer Centre, Ahmedabad for providing clinical specimens and assistance with clinical coordination. We acknowledge the National Facility for Gene Function in Health and Disease at IISER Pune for their invaluable support during the in vivo experimentation. We also thank Dr. Vishal Modi at Swasthyam Pathology Laboratory for their support in histopathological and immunohistochemical analysis. We acknowledge, Dept. of Bioinformatics, Gujarat University for providing computational resources for data analysis.

