## Supplementary material for "Inhibition of miR-1307 Reverses Resistance to Cisplatin in Drug-Resistant Oral Squamous Cell Carcinoma"

### Supplementary Figure 1:

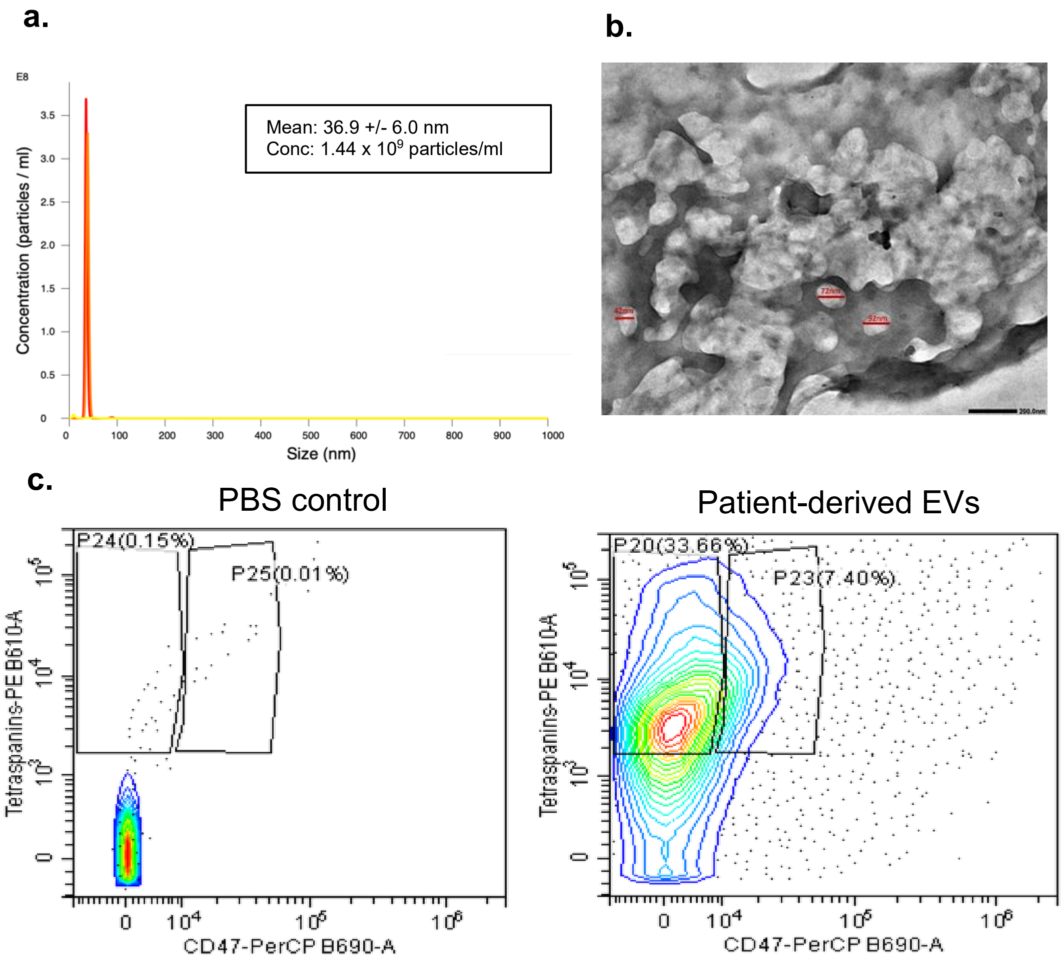


**Supplementary Figure 1: Characterisation of salivary EVs from OSCC patients**. (a) Estimation of concentration and size of salivary extracellular vesicles derived from OSCC patients using Nanoparticle Tracking Analysis (NTA). The Y-axis represents signal intensity and the X-axis indicates the size distribution of particles in NTA. It indicated an approximate concentration of $8.8\times10^6$ particles/mL (b) Representative TEM images of EVs from OSCC patients, revealed spherical morphology of the EVs, with a particle size distribution of 30–150 nm (Scale: 200 nm). (c) Flow cytometric analysis revealed strong expression of tetraspanins-PE (CD63, CD9, CD81) and CD47-PerCP in comparison to PBS control.

**Supplementary Figure 2:**

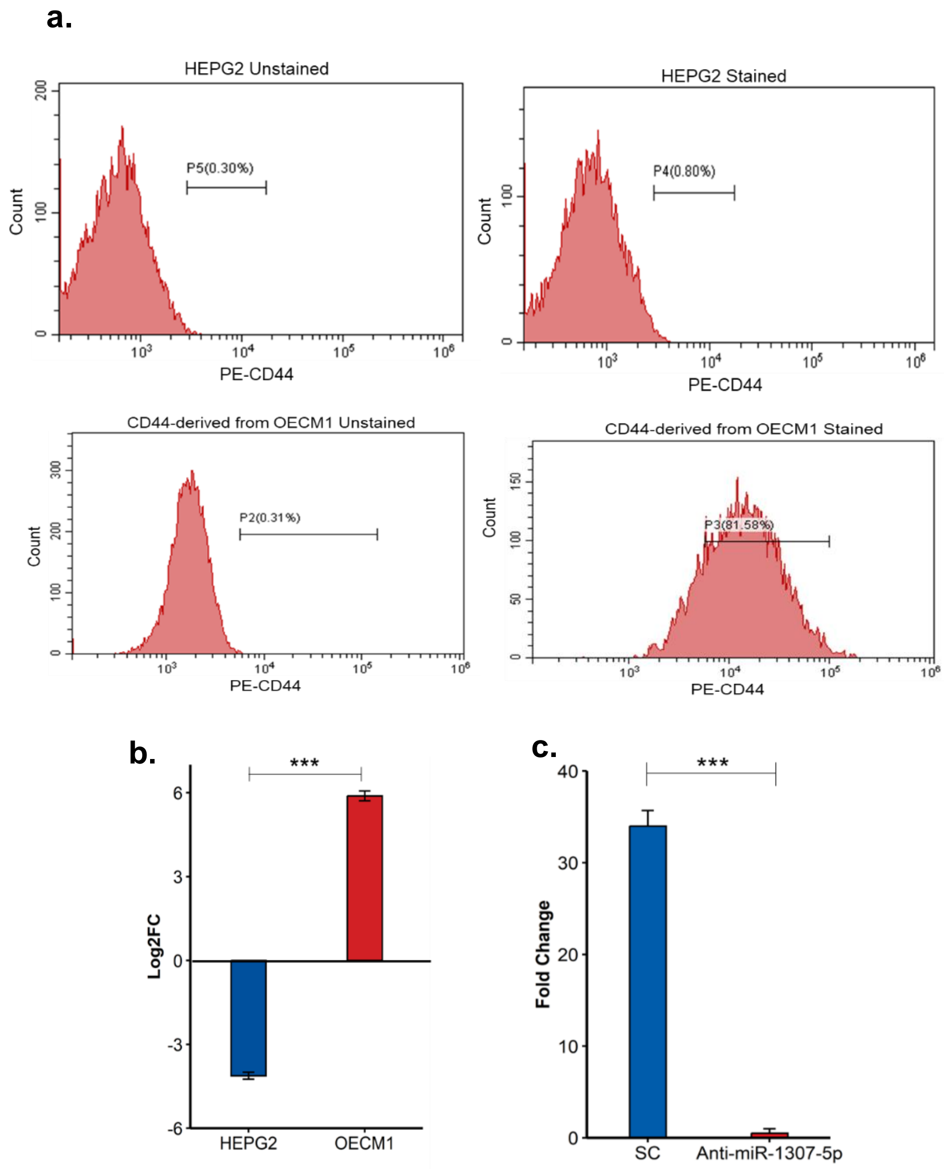


**Supplementary Figure 2: Characterisation of CD44 population in OECM1 cell line.** (a) Flow cytometry histogram showing the expression of CD44 detected using a PE-conjugated anti-CD44 antibody in the OECM1 cell line. HepG2 cells were used as a CD44-negative control. (b) qPCR analysis showing the relative expression of CD44 in the OECM1 and HEPG2 cell line.The data was normalised with B actin values, and relative expression of miRNA was analysed using the ΔΔCt method. (c) Representative bar-graph depicting miR-CD44 levels in CD44+ cells after transfection with anti-miR-1307-5p cells compared to the SC using real-time PCR. The data was normalised with U6 values, and relative expression of miRNA was analysed using the ΔΔCt method.

### Supplementary Figure 3:

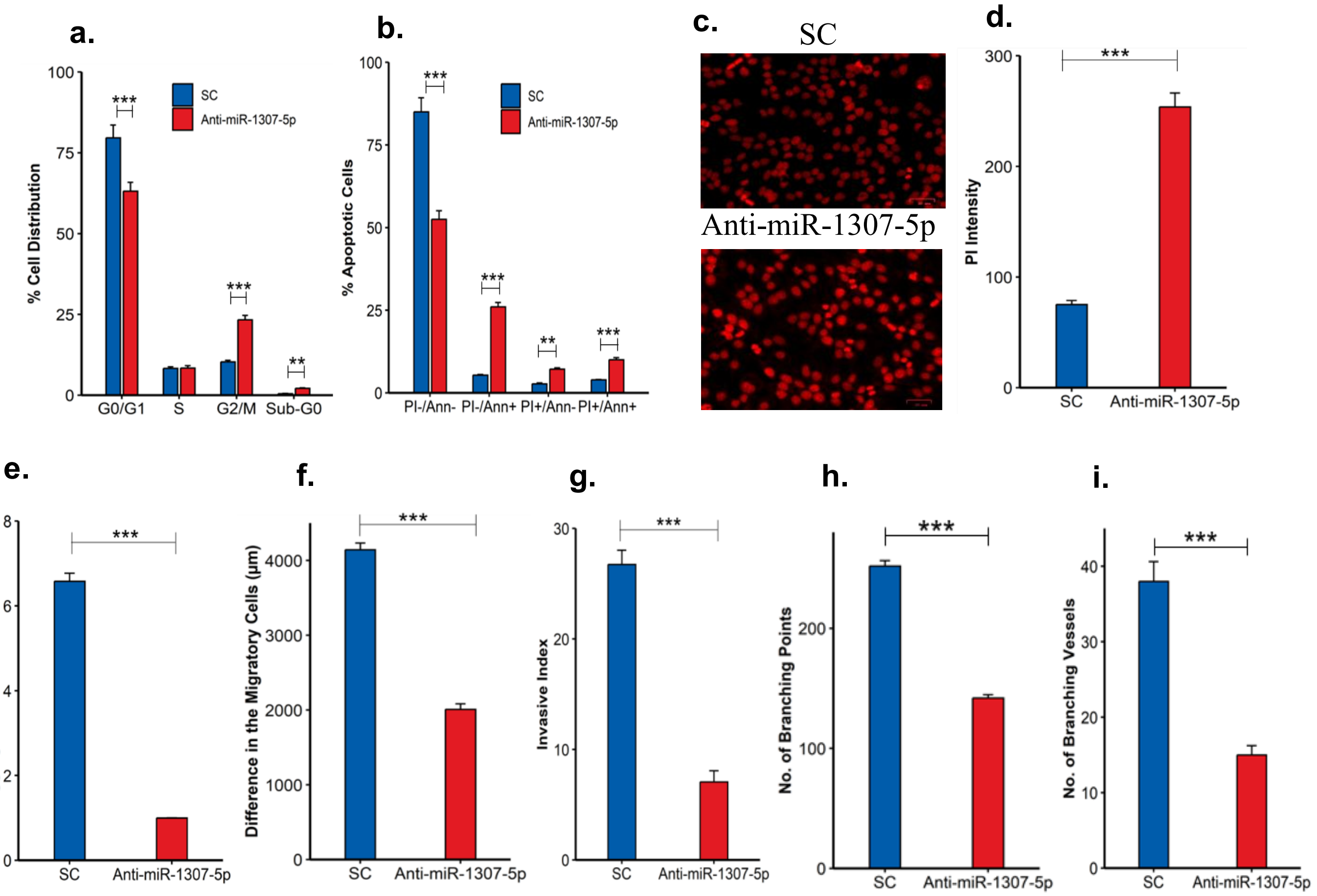


**Supplementary Figure 3: Downregulation of miR-1307-5p induces apoptosis and regulates epithelial-to-mesenchymal and angiogenic markers.** (a) Quantitative analysis of cell cycle distribution showed a significant increase in G2/M arrest and a corresponding decrease in G0/G1 in anti-miR-1307-5p compared to SC. (b) Quantitative analysis indicates a significant rise in early apoptotic cells (Annexin $V^+/PI^-$) in anti-miR-1307-5p-treated cells. (c) Fluorescence images of PI-stained $CD44^+$ cells (scale: 35 µm) showing enhanced apoptotic features—nuclear condensation and fragmentation—in the anti-miR-1307-5p group. (d) Quantitative bar graphs represent the fluorescence intensity of nuclear fragmentation and chromatin condensation in anti-miR-1307-5p-treated compared to SC. (e) Quantitative bar graphs represent the aggregate (red)-to-monomer (green) ratio in cells treated with anti-miR-1307-5p compared to SC. (f) Quantitative analysis of cell migration revealed a significant decrease in migratory potential in the anti-miR-1307-5p-treated cells compared to SC (g) Quantitative analysis of spheroid invasion shows a marked reduction in the invasiveness of anti-miR-1307-5p compared to SC. (h) Quantification of Calcein-AM fluorescence images of HUVEC tube formation in co-culture with treated $CD44^+$ cells, showing diminished endothelial network development in the anti-miR-1307-5p group. (i) IKOSA-based quantification of Ex-vivo CAM assay images showing reduced vascular branching and density after anti-miR-1307-5p treatment. Experiments were performed in triplicate, and error bars represent mean ± SD. The statistical significance is indicated as: (*$p < 0.05$; **$p < 0.01$; ***$p < 0.001$).

### Supplementary Figure 4:

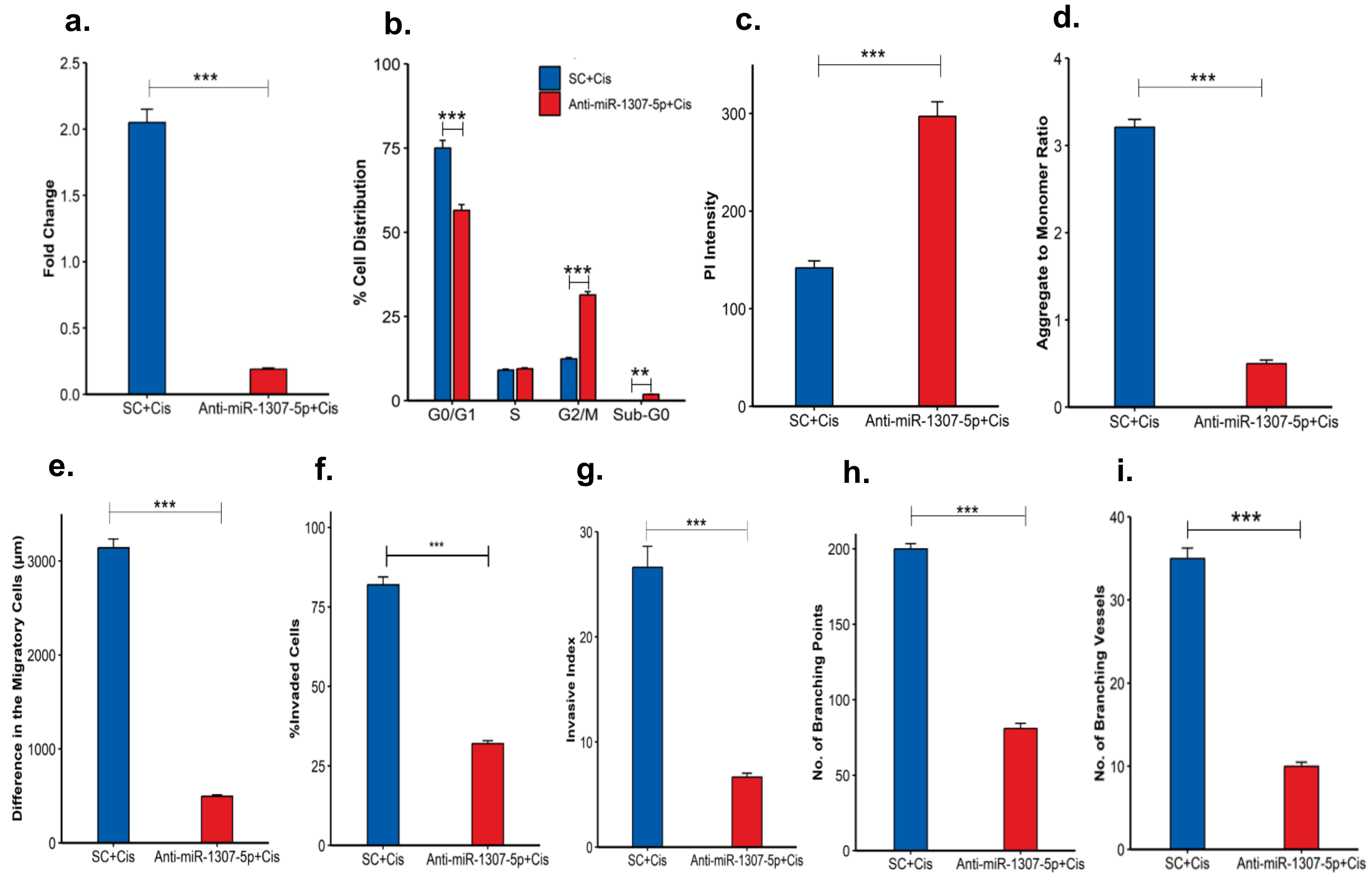


**Supplementary Figure 4: Anti-miR-1307-5p mediates chemoresistance in OSCC.** (a) Representative bar-graph depicting miR-1307-5p+Cis levels in CD44+ cells after transfection with anti-miR-1307-5p cells compared to the SC+Cis using real-time PCR. The data was normalised with U6 values, and relative expression of miRNA was analysed using the ΔΔCt method. (b) Quantitative analysis of PI-stained cells revealed that anti-miR-1307-5p+Cis induced G2/M population arrest accompanied by a decrease in G0/G1 relative to SC+Cis. (c) Quantitative bar graphs represent the fluorescence intensity of nuclear fragmentation and chromatin condensation in anti-miR-1307-5p+Cis-treated compared to SC+Cis (d) Quantitative bar graphs represent the red-to-green ratio in cells treated with anti-miR-1307-5p+Cis compared to SC+Cis. (e) Quantitative analysis of cell migration revealed a significant decrease in migratory potential in the anti-miR-1307-5p+Cis-treated cells compared to SC+Cis. (f) Transwell invasion assay quantification confirmed a marked reduction in the invasive capacity of $CD44^{+}$ CSCs following anti-miR-1307-5p+Cis treatment. (g) Quantitative analysis of spheroid invasion shows a marked reduction in the invasiveness of anti-miR-1307-5p+Cis compared to SC+Cis. (h) Quantitative analysis indicates a significant reduction in tube formation in anti-miR-1307-5p+Cis compared to SC+Cis. (i) Quantitative analysis with IKOSA-based quantification confirming significant reductions in vessel density and branching in the anti-miR-1307-5p+Cis-treated cells compared to SC+Cis. Experiments were performed in triplicate, and error bars represent mean ± SD. The statistical significance is indicated as: (*$p < 0.05$; **$p < 0.01$; ***$p < 0.001$).

### Supplementary Figure 5:

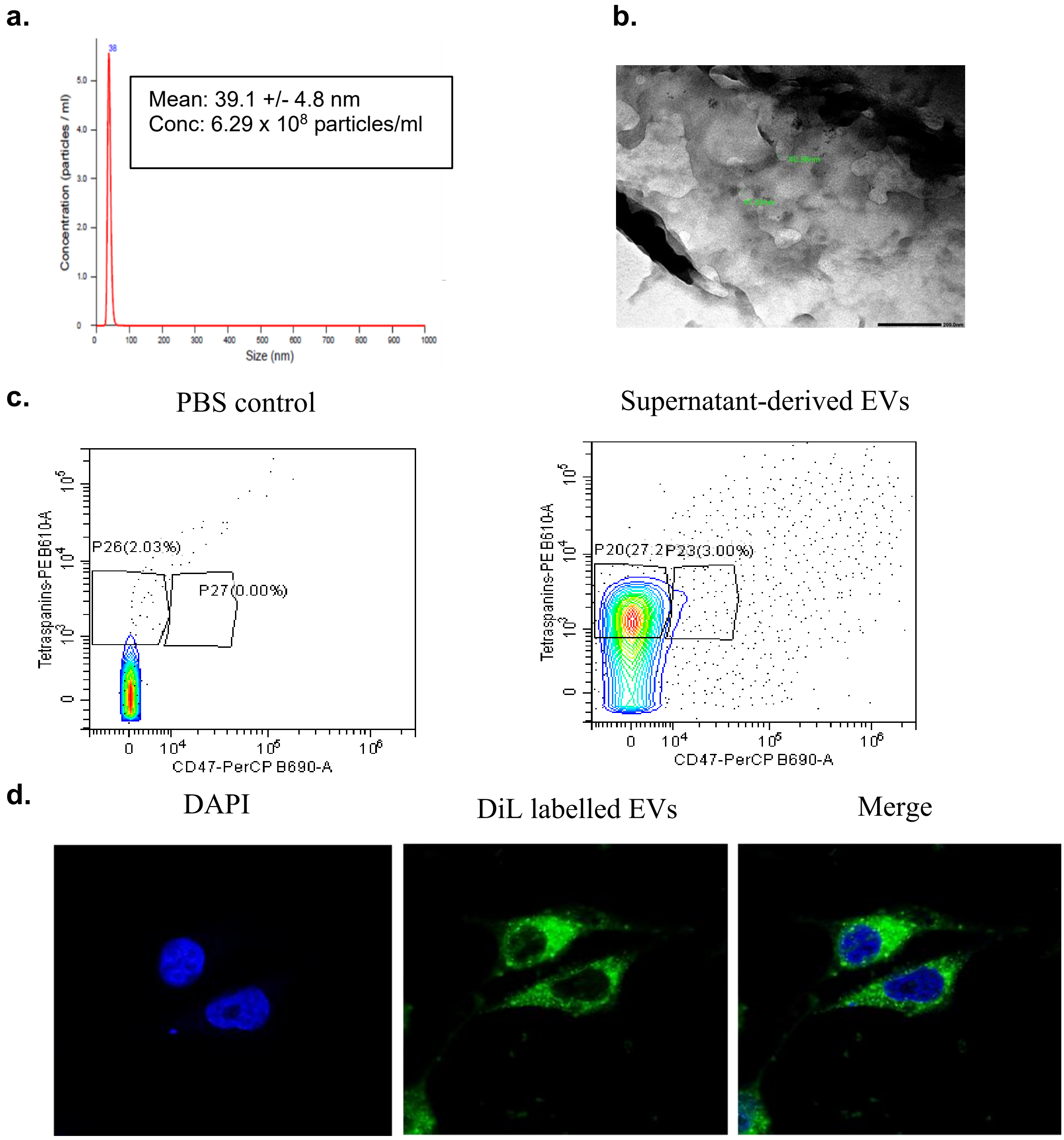


**Supplementary Figure 5: Characterisation of EVs isolated from CD44+ CSCs and abundance of anti-miR-1307-5p in EVs** (a) Characterisation of EVs derived from supernatant of CD44+ cells. Size: 32.9 nm and concentration: 8.1 × 10^7 particles/mL of EVs were determined using NTA. X-axis: size distribution of particles, Y-axis: signal intensity in the graph. (b) Representative TEM images of EVs representing a size of 40-50 nm. (c) The purity of EVs derived from supernatant of cells based on the surface markers was identified using flow cytometric analysis. EVs were stained with PE-conjugated antibodies against tetraspanins (CD63, CD81, and CD9) (d) Fluorescent cell analysis and representative confocal imaging for the uptake of CD44+-DiI-labelled EVs by recipient CD44+ cells (nuclei stained with DAPI). The data was normalised to SC, experiments were performed in triplicate, and error bars represent mean ± SD. The statistical significance is indicated as: (*p< 0.05; **p < 0.01; ***p < 0.001).

### Supplementary Figure 6:

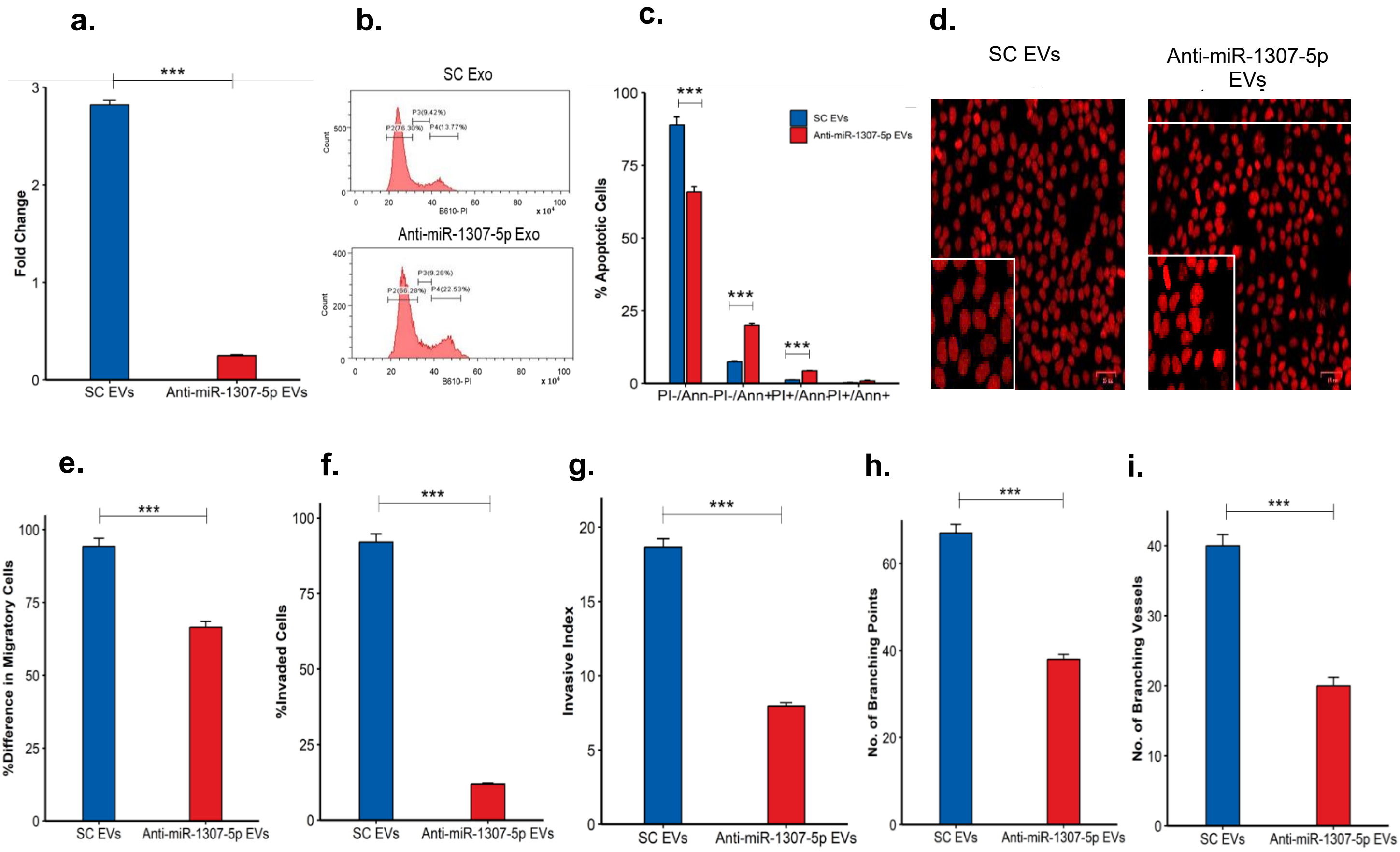


**Supplementary Figure 6- Anti-miR-1307-5p-EVs Impede Migration, Invasion, and Angiogenesis of CD44+ Cancer Stem Cells**

(a) Representative bar graph depicting miR-1307-5p expression in EVs derived from anti-miR-1307-5p-treated CD44+ cells compared to the scramble control (SC), as quantified by real-time PCR. The data were normalized to β-actin, and relative expression levels were calculated using the ΔΔCt method (b) Flow-cytometry histograms of PI-stained DNA content, showing a significant increase in G2/M arrest and a decrease in G0/G1 in anti-miR-1307-5p EVs compared to SC EVs. X-axis: PI fluorescence at 610nm wavelength, Y-axis: cell-count distribution. (c) Representative bar graph showing a higher percentage of early apoptotic (Annexin-V-FITC+/PI-) and dead (Annexin-V-FITC+/PI+) cells in CD44+ cells treated with anti-miR-1307-5p EVs compared to SC. (d) Fluorescent microscopic images of CD44+ cells treated with anti-miR-1307-5p EVs or SC EVs (scale: 35 µm). Nuclear fragmentation and chromatin condensation indicate the formation of apoptotic bodies, suggesting the induction of apoptosis in cells treated with anti-miR-1307-5p loaded EVs (e) The bar graph depicting the reduced migration of cells treated with anti-miR-1307-5p-EVs (scale bar: 10 µm) after 48 hours (f) Representative bar graph for the transwell assay demonstrating a significant decrease in the proportion of invaded cells when treated with anti-miR-1307-5p, EVs as opposed to SC EVs (g) Quantitative analysis of spheroid invasion shows a marked reduction in the invasiveness of anti-miR-1307-5p compared to SC respectively. (h) The representative bar graph shows the change in branching points, illustrating the reduced angiogenic potential in the anti-miR-1307-5p EV-treated group compared to the SC EV-treated group (i) Representative bar graph quantifies the difference in branching points, demonstrating reduced angiogenesis in anti-miR-1307-5p EV-treated CAMs. All data are presented as the mean ± SE of three independent experiments compared to the respective scramble control. Error bars represent mean ± SD of three independent experiments. The statistical significance is indicated as: (*$p < 0.05$; **$p < 0.01$; ***$p < 0.001$).

**Supplementary Table 1**

**PRIMER SEQUENCES**

| miRNA | Primer Sequences |
|---|---|
| U6 | F 5’-CGGGCTCGACCGGACCTCG-’3<br>R 3’-CAGCCACAAAAGAGCACAAT-5’ |
| 1307-5p | F 5’-GGAAACAGCTATGACCATG-3’<br>R 3'-GGTTTTCCCAGTCACGAC-5’ |

| Gene | Primer Sequences |
|---|---|
| CD44 | F 5'-CGGACACCATGGACAAGTTT-3'<br>R 5'-CCGTCCGAGAGATGCTGTAG-3' |
| VEGF-α | F 5'-GCTGTCTTGGGTGCATTGG-3'<br>R 5'-GCAGCCTGGGACCACTTG-3' |
| N-CADHERIN | F 5’-CCACGCCGAGCCCCAGTATC-3’<br>R 5’-CCCCCAGTCGTTCAGGTAATCA-3’ |
| E-CADHERIN | F 5’-ATTCTGATTCTGCTGCTCTTG-3’<br>R 5’-AGTCCTGGTCCTCTTCTCC-3’ |
| PI3K | F 5’-ACCACTACCGGAATGAATCTCT-3’<br>F 5’-GGGATGTGCGGGTATATTCTTC-3’ |
| AKT | F 5’-ACTCATTCCAGACCCACGAC-3’<br>F 5’-AGCCCGAAGTCCGTTATCTT-3’ |
| YAP1 | F 5’-CTCAAAATCCAGTGTCTTCTCC-3’<br>F 5’-TCACCTGTATCCATCTCATCC-3’ |
| VIM | F 5’-CTCTCAAAGATGCCCAGGAG-3’<br>F 5’-GCACGATCCAACTCTTCCTC-3’ |
| ß -Actin | F 5’ -CATGTACGTTGCTATCCAGGC -3’<br>R 5’ -CTCCTTAATGTCACGCACGAT -3’ |

Table 1- The table represents the primer sequences of the miRNA and genes.
